# Experimental evolution reveals the synergistic genomic mechanisms of adaptation to ocean warming and acidification in a marine copepod

**DOI:** 10.1101/2021.11.01.466754

**Authors:** Reid S. Brennan, James A. deMayo, Hans G. Dam, Michael Finiguerra, Hannes Baumann, Vince Buffalo, Melissa H. Pespeni

## Abstract

Metazoan adaptation to global change will rely on selection of standing genetic variation. Determining the extent to which this variation exists in natural populations, particularly for responses to simultaneous stressors, is therefore essential to make accurate predictions for persistence in future conditions. Here, we identify the genetic variation enabling the copepod *Acartia tonsa* to adapt to experimental ocean warming, acidification, and combined ocean warming and acidification (OWA) conditions over 25 generations. Replicate populations showed a strong and consistent polygenic response to each condition, targeting an array of adaptive mechanisms including cellular homeostasis, development, and stress response. We used a genome-wide covariance approach to partition the genomic changes into selection, drift, and lab adaptation and found that the majority of allele frequency change in warming (56%) and OWA (63%) was driven by selection but acidification was dominated by drift (66%). OWA and warming shared 37% of their response to selection but OWA and acidification shared just 1%. Accounting for lab adaptation was essential for not inflating a shared response to selection between all treatments. Finally, the mechanisms of adaptation in the multiple-stressor OWA conditions were not an additive product of warming and acidification, but rather a synergistic response where 47% of the allelic responses to selection were unique. These results are among the first to disentangle how the genomic targets of selection differ between single and multiple stressors and to demonstrate the complexity that non-additive multiple stressors will contribute to attempts to predict adaptive responses to complex environments.

## Introduction

Human activity is driving dramatic environmental changes across the globe. A crucial challenge in evolutionary biology is understanding how adaptation will proceed under this rapid change. After rapid environmental shifts, populations may go extinct unless phenotypes evolve through selection on standing genetic variation already present in the population. It is established that selection due to environmental shifts can act on this standing variation (Barrett and Schluter, 2008) and can result in rapid phenotypic (Bumpus, 1899; Donihue et al., 2018) and genomic changes (Campbell-Staton et al., 2017; De Wit et al., 2014; Rodríguez-Trelles et al., 2013). Much effort has focused on identifying the genes that underlie adaptive genetic variation in natural populations using gene-environment-associations or quantitative genetic methods, including genome-wide association studies (Waldvogel et al., 2020).

An alternative framework is evolve-and-resequence (E&R), where populations are evolved under a selective environment and sequenced at various time points (Schlötterer et al., 2015). Allele frequency changes can then be analyzed to find the loci under selection and to understand the genetic architecture of the adaptive response (Castro et al., 2019). This method is powerful in that it reveals both the presence of standing adaptive genetic variation and the mechanistic basis of adaptation at the genetic level. While the majority of E&R has been pioneered in the *Drosophila* model system, the approach has since expanded to other species, including marine organisms (Kelly and Griffiths, 2021). However, disentangling the effects of drift and selection on allele frequency changes remains a formidable challenge for E&R studies (Vlachos et al., 2019). This is further complicated when adaptation proceeds from standing genetic variation on polygenic traits, which can lead to diffuse and subtle signals of selection that can be difficult to distinguish from drift (Kemper et al., 2014; Pritchard et al., 2010). New covariance-based methods can distinguish selection from drift using replicated genomic samples through time (Buffalo and Coop, 2020, 2019). Nevertheless, incorporating appropriate experimental controls is still essential to prevent spurious results from phenomena such as adaptation to laboratory (lab) conditions or drift (Buffalo and Coop, 2020; Castro et al., 2019).

The world’s oceans are particularly vulnerable to human activity as unprecedented increases in atmospheric CO2 lead not only to higher global temperatures, but also decrease ocean pH due to CO2 dissolution, a process called ocean acidification (Doney et al., 2009; Pelejero et al., 2010). These environmental changes could fundamentally alter marine ecosystems, particularly if they affect zooplankton that link primary producers with higher level consumers and are essential in ocean biogeochemical cycles (Banse, 1995; Dam et al., 1995; Möllmann et al., 2008). For zooplankton, pH declines may increase the cost of acid-base regulation (Thor and Dupont, 2015) while temperature exposure beyond an organism’s optimum drives quick performance declines (Sasaki and Dam, 2021). As such, there has been considerable effort to understand how zooplankton will be able to evolve in response to shifting global conditions (De Wit et al., 2016; Griffiths et al., 2021; Kelly et al., 2017, 2012; Langer et al., 2019; Lee et al., 2011, 2007). However, a limitation of previous work measuring evolutionary potential has been the focus on single stressor selective pressures. Given the simultaneous shift in abiotic factors occurring under climate change, it is essential to understand the combined effects of multiple stressors (Boyd et al., 2018). This is especially important as multiple stressors can have additive, antagonistic, or synergistic effects (Gunderson et al., 2016) and studying stressors both alone and in combination can help reveal the physiological and evolutionary processes that promote resilience.

*Acartia tonsa* is one of the most abundant copepods globally and a dominant species in coastal ecosystems where it is a primary food source of numerous fish (Turner, 1984). Given this, the resilience of *A. tonsa* is essential in preserving ecosystem functioning under global change. We previously conducted a selection experiment where we reared *A. tonsa* for 25 generations (^~^1 year) to characterize fitness-related trait responses to warming, acidification, and their combination (OWA)(Dam et al., 2021). We found that exposure to combined warming and acidification conditions (OWA) drove a decrease in population fitness that largely recovered by three generations through improved hatching success and earlier reproductive timing, revealing rapid but incomplete adaptation of *A. tonsa* to OWA selection. In contrast, warming and acidification alone had minor effects on population fitness.

Here, we use the same populations from our evolution experiment to measure genomic responses to 25 generations of selection in ambient, warming, acidification, and OWA conditions; asking specifically: 1) What is the relative role of selection versus drift in driving evolutionary change? 2) To what degree are adaptive changes shared between the single and multiple stressors? 3) What are the genes and functional mechanisms underlying rapid adaptation to warming, acidification, and OWA conditions? We demonstrate that *A. tonsa* has sufficient adaptive potential to evolve in response to warming, acidification, and OWA conditions, show the large effects of warming on copepod adaptation relative to acidification, and reveal the synergistic effects of warming and acidification when combined as a multiple stressor. Finally, these results reveal the importance of controlling for lab adaptation in E&R experiments.

## Results

### Genome-wide variation in response to experimental selection

To quantify genomic response to selection of *Acartia tonsa*, samples were collected from each of four replicates of acidification, warming, and OWA treatments at generation 25 and from an ambient control at both generation 0 and 25. We designed 32,413 capture probes to target both regulatory (11,102) and coding (21,311) regions of the genome and used pooled-capture sequencing of 50 individuals per replicate to characterize the allele frequency changes across the experiment. After filtering, we obtained 394,667 single nucleotide polymorphisms (SNPs) with no missing data across all samples. The founding population possessed high genome-wide genetic diversity on which selection could act (Tajima’s π: 0.0148 ± 0.0111). Principal component analysis PCA (Fig. 1) showed that the variance in genome-wide allele frequencies for all samples clustered by treatment group, and F25 samples had diverged from the F0 founding population along principal component 1 (15.82% of the variation). There was also separation of the warming and OWA treatments from ambient and acidification treatments along this axis. Principal component 2 (10.99% of the variation) further separated each treatment group into distinct clusters.

**Figure 1:**
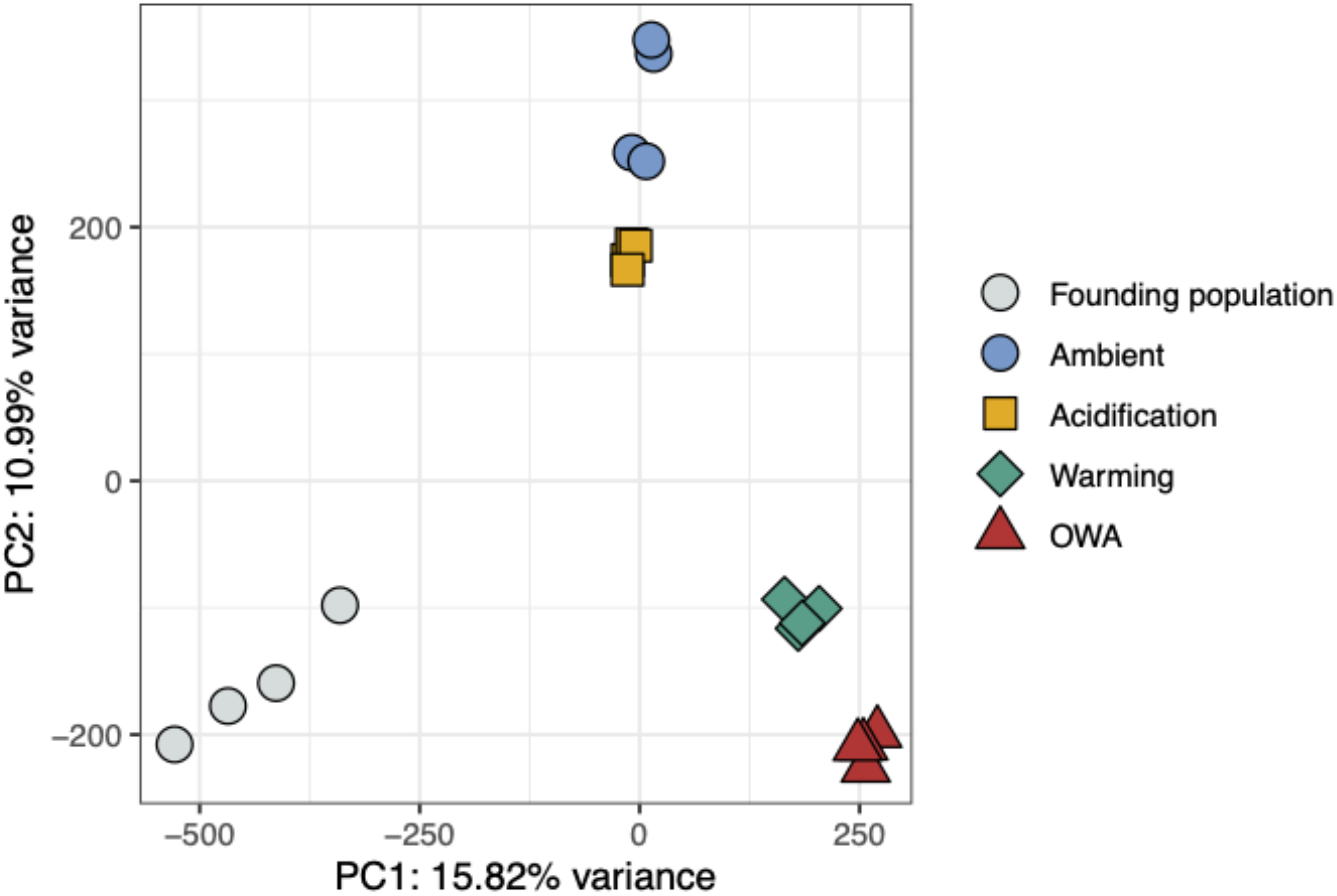
Principal component analysis of allele frequencies from 394,667 loci across the genome where color and shape distinguish treatment groups. The founding population is F0, all others are from F25.

### Contribution of drift, selection and lab adaptation to allele frequency change

Leveraging both the replicated and temporal nature of our experimental design, we used the covariance-based method, *cvtk* (Buffalo and Coop (2020), to identify even subtle contributions of selection to genome-wide allele frequency change. We calculated pairwise covariances in allele frequency change from F0 to F25 in 10,000 base pair windows between all pairwise replicates (Fig. S1). From this, we determined the convergent correlation, a standardized measure of how similar allele frequency changes are between replicates or different treatments due to a convergent selection response (Fig. 2A). Under drift alone, allele frequency changes would be inconsistent across replicates, leading to a convergent correlation of zero. By contrast, populations could have parallel changes in allele frequency due to a shared response to a selective pressure, leading to a high convergent correlation. As expected, we observed the highest convergent correlations between replicate pairs within a selection regime, indicative of parallel responses to selection pressure (Fig. 2A). By contrast, comparisons between replicates across treatment groups generally had weaker convergent correlations than the intra-treatment comparisons (Fig. 2A; see Fig. S12 for 95% bootstrapped confidence intervals). The exception to this was between OWA and warming treatments where the inter-treatment convergent correlations approached or exceeded within treatment comparisons (Fig. 2A). This high similarity in response to selection between OWA and warming indicates shared targets of selection between the two conditions; this pattern was not present between other inter-treatment comparisons.

**Figure 2:**
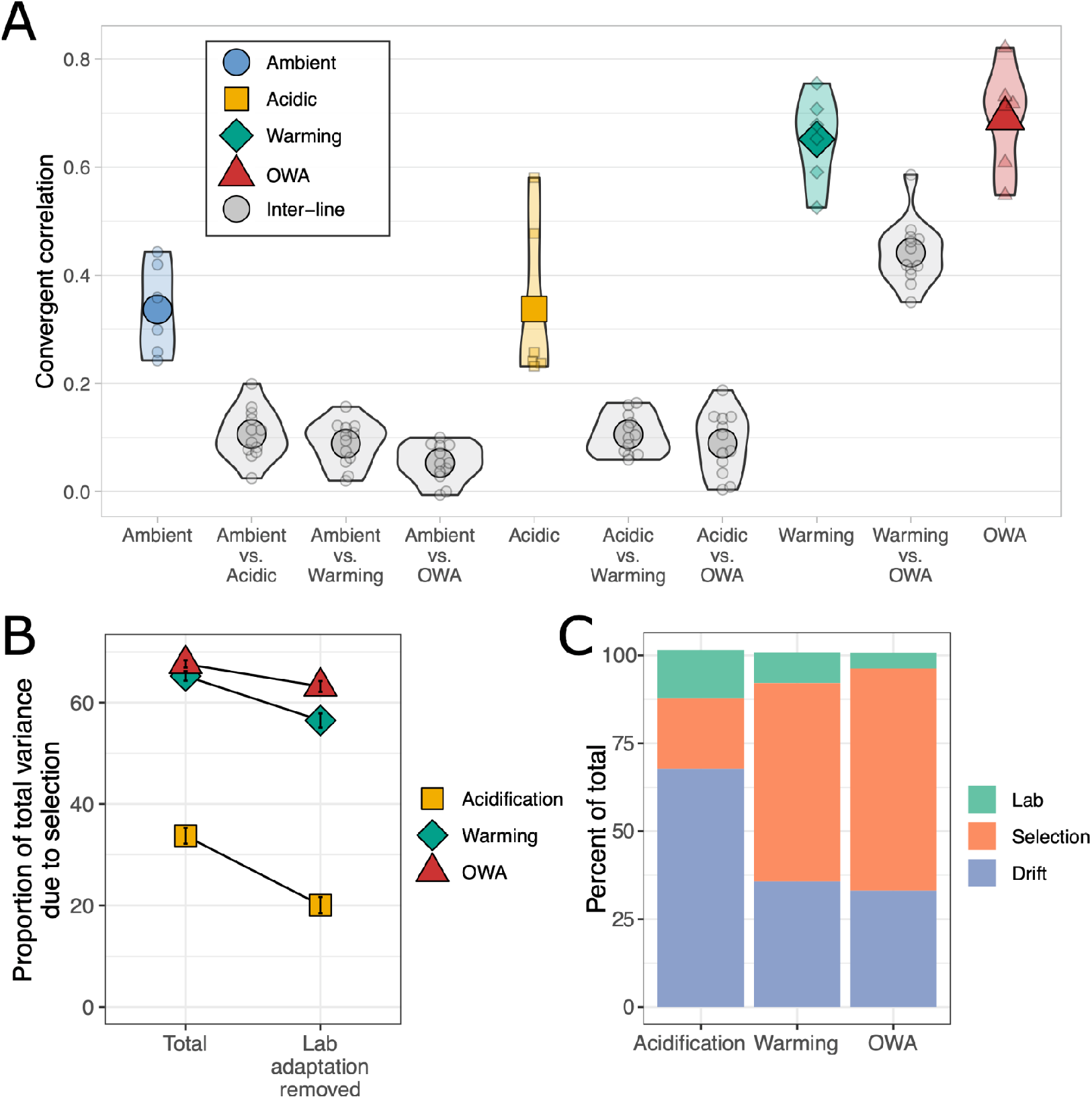
A) Convergent correlation of allele frequency change from F0 to F25 between treatment groups. Higher values indicate a more similar change in allele frequency between two samples. Each small point is the convergent correlation between two samples and the large points represent the mean of each group. For visual clarity we omit the confidence intervals of each pairwise convergent correlation, see Fig. S12 for these values. B) Proportion of total variance in allele frequency due to selection with and without accounting for lab adaptation. Lab adaptation was removed by quantifying the shared response between the ambient and each treatment group. Error bars are 95% bootstrapped confidence intervals. C) The contribution of lab selection, treatment selection, and drift to the total variance of allele frequency change from F0 to F25. The lab component was determined using the covariance of allele frequency change between the ambient and each treatment group while selection was identified as the covariance within a treatment group minus the lab adaptation. The remaining variance was attributed to drift.

Within treatments, we found that OWA had the highest variance in allele frequency change (0.0536, 95% CI[0.0521, 0.0551]), followed by warming (0.0410, CI[0.0399, 0.0422]) and acidification (0.0243, CI[0.0237, 0.0250]); higher values indicate larger and more abundant allele frequency shifts across the genome. The contribution of selection to this variance in allele frequency change was then quantified by calculating the shared changes, as pairwise covariances, between all replicates within a treatment (Fig. 2B). The remaining variance not due to selection was attributed to drift or unique selection responses between the replicates, which we could not distinguish here (Barghi et al., 2019). The OWA line was most strongly affected by selection with 67.6% (CI[66.9, 68.3]) of the total variance in allele frequency change due to selection. Warming was slightly, but significantly, lower (65.2% CI[64.3, 66.2]) than OWA, and acidification showed the lowest percentage of variance due to selection at 33.7% (CI[32.2, 35.3]).

The movement of the ambient lines in principal component space relative to the founding populations (Fig. 1) as well as the relatively high covariance between these samples (Fig. S1) suggested adaptation to laboratory (lab) conditions unrelated to treatments. Thus, estimates of shared variance between samples could be inflated due to the shared variance driven by lab selection. Because the ambient and any treatment line shared only the lab environment, but not the selective pressure of interest, the covariance between ambient and selected line represented adaptation to the lab environment and could therefore be estimated and accounted for. The total estimated lab effects were lower in OWA conditions (0.0024, 95% CI[0.0018, 0.0029]) than either warming (0.0036, CI[0.003, 0.0041]) or acidification (0.0033, CI[0.0029, 0.0038]). After accounting for the lab adaptation signal, the percent of total allele frequency change variance due to selection decreased for all treatments, but the rank order and significance remained: OWA 63.2% CI[62.1, 64.2], warming 56.5% CI[55.1, 57.9], acidification 20.1% CI[18.5, 21.7]. The drift component was much larger for acidification than warming or OWA (Fig. 2C).

Last, we quantified the shared response to selection between OWA, warming, and acidification with and without removing estimated lab adaptation effects. We calculated the mean pairwise covariance between each group scaled by the total variance. When lab adaptation was not removed, acidification shared a response with both OWA (8.4% CI[7.0, 9.8]) and warming (10.3% CI[8.9, 11.8], Fig. 3). However, when lab adaptation was removed, acidification shared no response with warming (−0.25% CI[-1.36, 0.87]) and a minimal response with OWA (1.12%, CI[0.02, 2.21]). In contrast, the OWA and warming lines shared a large proportion of their allele frequency change both with (37.1% CI[35.9, 38.4]) and without (43.5% CI[42.3, 44.6]) accounting for lab adaptation.

**Figure 3:**
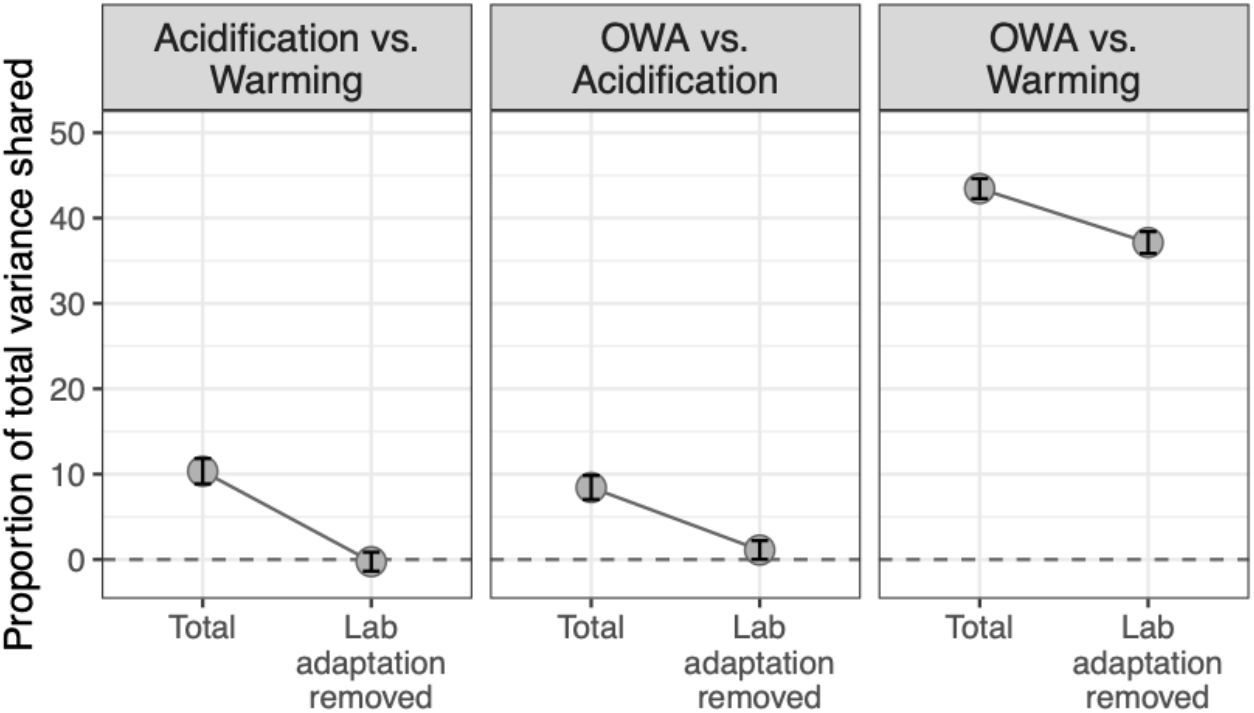
Shared response to selection between treatments with and without accounting for lab adaptation. Error bars represent 95% bootstrap confidence intervals.

### Identifying the specific targets of selection

While the covariance-based method above disentangled the contributions of selection and drift to the total variance in allele frequency change, it did not identify the specific genomic regions and loci likely underlying adaptation. To reveal the genetic variation responding to selection rather than drift, we simulated the degree of drift expected with our experimental design across 25 generations (see methods) and used these simulations with Cochran–Mantel–Haenszel (CMH) tests to identify loci that were evolving more than we would expect by drift alone in each treatment. We found 23,759 (6.0% of all SNPs), 18,324 (4.6%), and 7,516 (1.9%) SNPs that were candidate targets of selection for OWA, warming, and acidification, respectively (Figs. 4, S2). The number of candidate SNPs identified here likely represents an overestimation of the true targets of selection due to linkage; our simulations assumed independence between loci, which is not accurate as physical linkage is present in these data but decays to background levels after ~200 basepairs (Fig. S3). We again saw evidence for lab adaptation with 12,126 candidate SNPs (3.1% of total SNPs) exceeding the null drift expectations in the ambient lines. To control for this, we removed candidate SNPs from all treatments that were identified as evolving in the ambient treatment. The lab adaptation response was highly overlapping across the three treatment groups, with 2,940 ambient (i.e., lab adaptation) SNPs found in all three groups (Fig. S4). After removing any lab adaptation loci, we found 19,097 (4.8%), 13,309 (3.4%), and 3,400 (0.9%) candidate SNPs in OWA, warming, and acidification, respectively (Fig. 4). Comparing sets of candidate SNPs between treatments, OWA and warming had greater fold enrichment (FE) in overlapping candidate SNPs than expected by chance (observed: 6359; expected: 644; FE: 9.9; P < 0.0001) as did OWA and acidification (observed: 886; expected: 165; FE: 5.39; P < 0.0001), though to a lesser extent, indicating a greater degree of shared response to selection between OWA and warming than OWA and acidification (Fig. 4).

**Figure 4:**
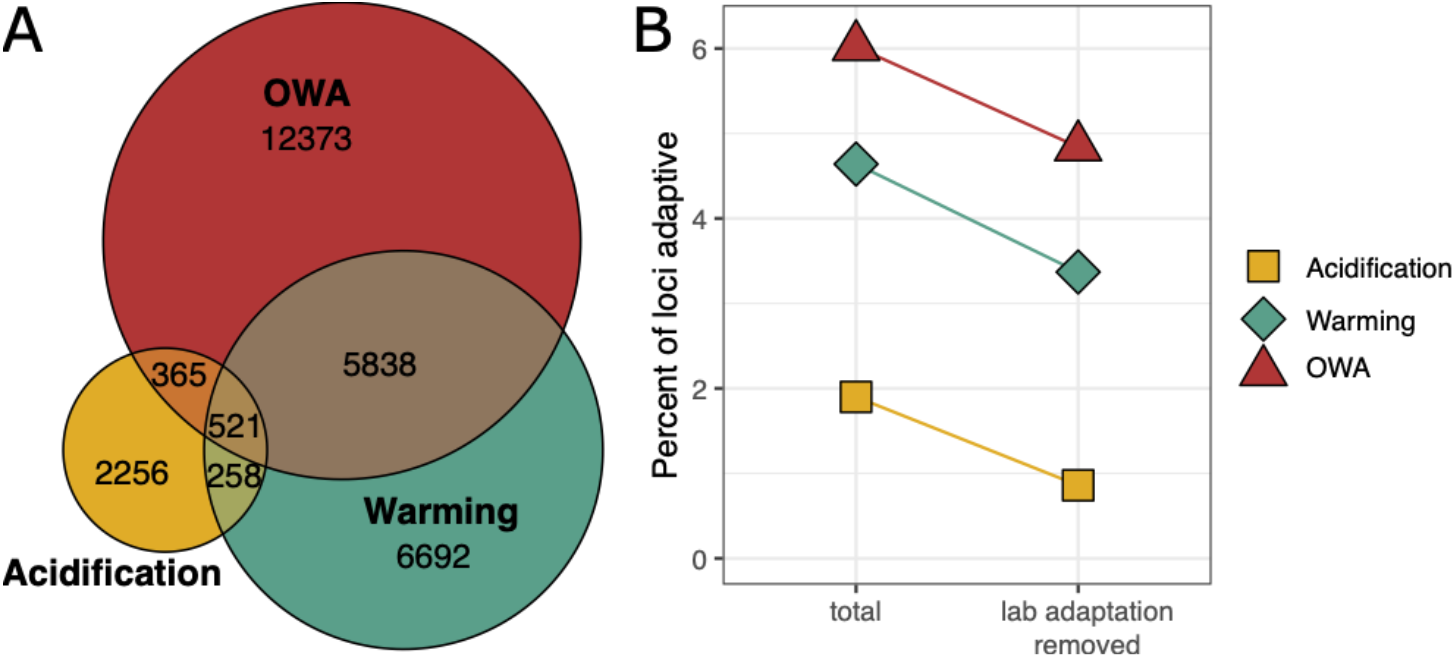
A) Overlap in candidate SNPs exceeding drift expectations identified from Wright-Fisher simulations with lab adaptation loci removed. The overlap between OWA and warming showed greater enrichment in their overlap (9.9 fold enrichment) than OWA and acidification (5.4 fold enrichment). B) Percent of total polymorphic loci identified as candidates under selection.

To further characterize the genomic regions responsive to selection, we tested for overrepresentation of candidate loci in coding vs. promoter regions and in annotated vs. unannotated gene regions. The candidate loci were unequally distributed across the genome and were overrepresented in coding regions for OWA (12,691 OWA candidate loci in coding regions/19,097 all OWA candidate loci = 0.665) and warming treatments (8,922/13,309 = 0.670) relative to genome-wide patterns (229,898 loci in coding regions /394,667 all loci = 0.583) (Table S1; Chi-squared test P < 0.001). Additionally, for all three treatments, there were fewer candidate loci on scaffolds with no gene annotations, suggestive of regions that were fragmented or non-functional (P < 0.001; genome-wide: 54,228/394,667 = 0.137; OWA: 1,384/19,097 = 0.072; warming: 908/13,309 = 0.069; acidification: 339/3,400 = 0.010). Together, the inflation of candidate loci in coding regions of annotated genes suggests that selection was disproportionately targeting protein coding variation during the adaptation process.

Our previous results showed that animals in the multiple-stressor OWA conditions recovered fitness, population growth rate, after just three generations (Dam et al., 2021). To determine the extent to which adaptive signals at F25 were also under selection and potentially driving phenotypic responses at F3, we quantified alleles identified as adaptive to OWA conditions at F25 that had also evolved in the adaptive direction after three generations in OWA conditions. We identified the loci with the top 0.1% significance values for OWA conditions at F25 and quantified the direction of allele frequency change of these same loci at F3 relative to F0. At F3, 72% of these top 370 most significant loci had shifted frequency in the same direction as F25 (Fig. 5A), exceeding the neutral expectation (P= 0.0008; Fig. 5B). This pattern held when the significance threshold was expanded to the top 1% and 10%. Additionally, F3 samples in PCA space were intermediate to F0 and F25 samples (Fig. S5). Together, these results suggest that selection was driving an adaptive response after only three generations in OWA conditions.

**Figure 5:**
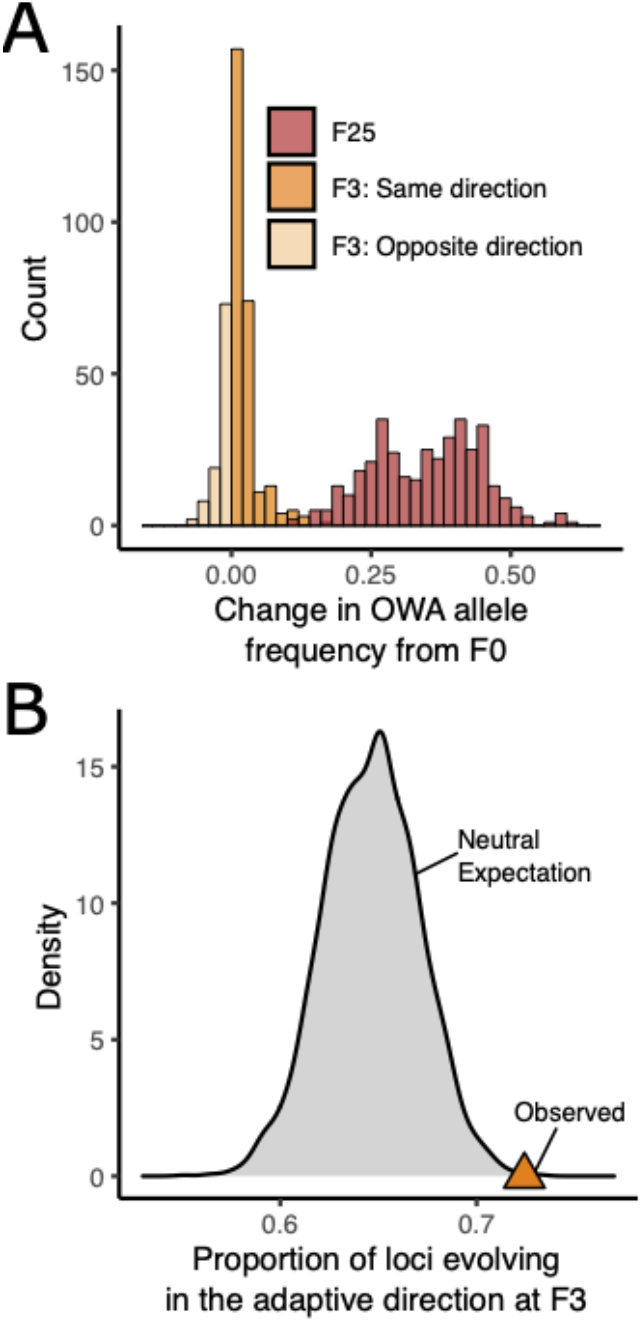
Allele frequency change of top 0.001 most significant loci (370 loci) in the OWA treatment at F3 and F25. A) For generation F3, color indicates if the locus is evolving in the same or opposite direction of the locus at F25. B) Neutral expectation of the proportion of loci that would be evolving in the same, adaptive direction at F3 as at F25. The neutral expectation was generated by randomly sampling 370 loci and calculating the proportion changing frequency in the same direction at F3 and F25. The observed data are more extreme than the neutral expectation (P= 0.0008).

### Functional targets of selection

Tests for gene ontology (GO) functional enrichment were used to gain insight into the physiological mechanisms that may be underlying adaptation to the different conditions. We tested for enrichment within the candidate SNPs with lab adaptation removed (Figs. 6, S8, Table S2). There was shared enrichment across all treatments for related processes corresponding to response to stimulus, cellular homeostasis, and stability, consistent with a general stress response and evolution for homeostasis. Warming and OWA were enriched for processes related to development while OWA and acidification were enriched for cell cycle processes, indicating shared targets of selection between OWA and each condition. There were also unique responses related to resilience: warming was enriched for response to heat, metabolic processes, and hypoxia, while acidification was enriched for protein transport and refolding, cytoskeleton and actin regulation, and gene expression regulation. Finally, the OWA line was enriched for processes related to ion transport. We also identified candidate genes, Na^+^,K^+^-ATPase subunit α-4, focal adhesion kinase 1, NADH dehydrogenase 1 alpha and beta, and DnaJ heat shock protein family (Hsp40) member A3, which had multiple loci responding to selection and were involved in the GO enrichment (Fig. S9). For these genes, linked haplotypes were increasing in frequency across hundreds of base pairs; the extent of linkage we could observe was limited by the number of capture probes within each gene, generally one.

**Figure 6:**
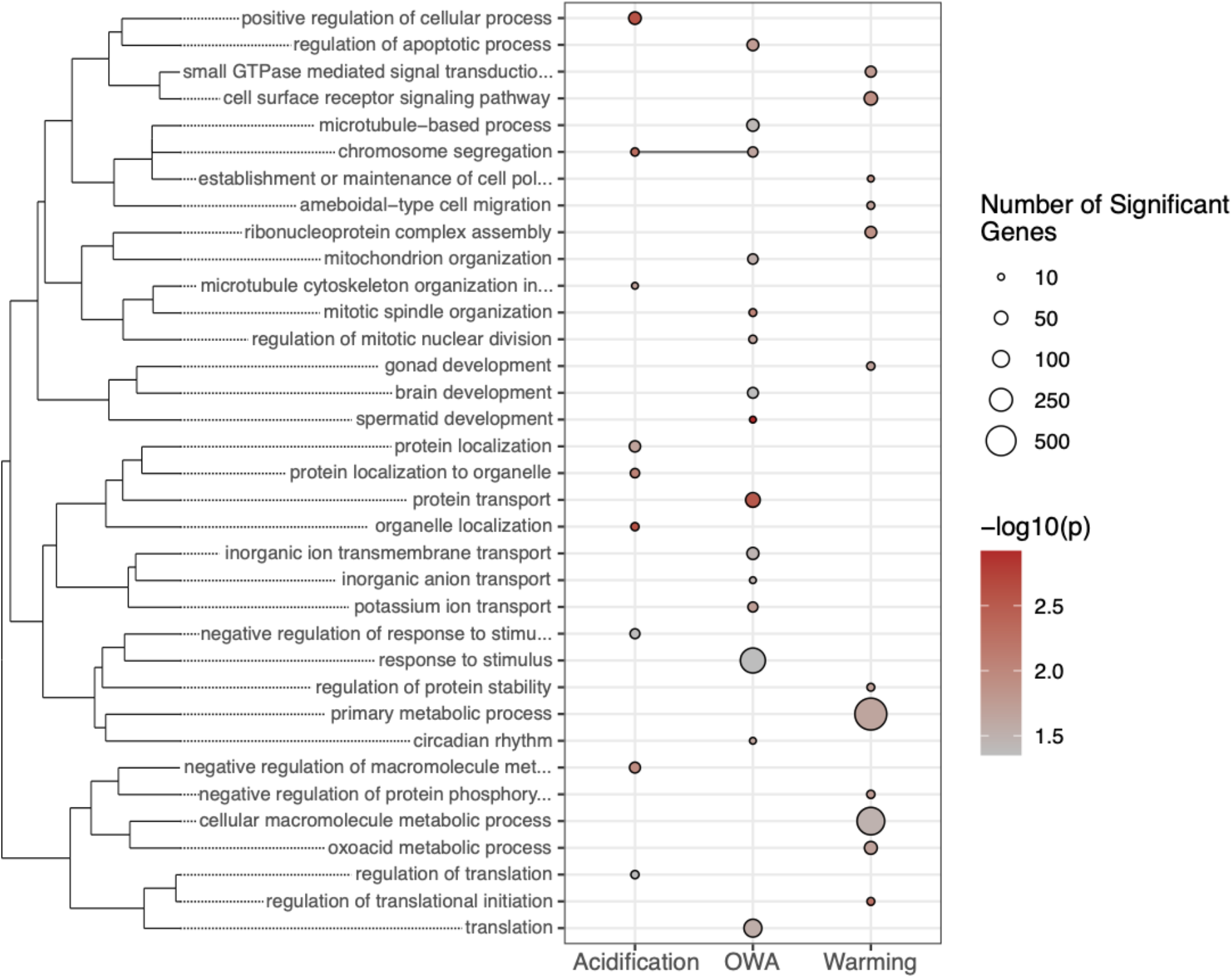
Gene ontology (GO) enrichment from candidate SNPs under selection. The size of a point indicates the number of genes in a category and color shows p-value. GO terms are clustered by similarity and the presence of a dot in a column indicates significant enrichment for that term in the treatment condition. For clarity, only GO terms with at least 10 significant genes are shown. See fig. S8 for full results.

### Polygenic soft sweeps from standing genetic variation

When selection acts on standing genetic variation in natural populations, one expects adaptive variation to be present in multiple individuals and spread across genetic backgrounds, which leads to soft sweeps. In addition, when a trait is polygenic, selection can result in the simultaneous, subtle, shift of many loci across the genome. As such, the dramatic bottlenecks and signals of selection that are common under hard sweeps are not expected (Bernatchez, 2016; Messer and Petrov, 2013; Pritchard and Di Rienzo, 2010). We found evidence that adaptation was proceeding through soft sweeps of polygenic traits. In addition to the large number of loci across the genome responding to selection (Fig. 4), we found only modest increases in linkage disequilibrium that scaled with the strength of selection where the OWA lines showed the largest increase relative to the founding population (Fig. S3), suggesting that hard sweeps were not present. Similarly, we found no link between the response to selection and genome-wide genetic diversity (Tajima’s π), though diversity fell by 7-14% across all treatment groups from the starting diversity of 0.0148 ± 0.0111 (Fig. S6; Wilcoxon signed-rank test, P < 0.0001). This drop was greatest in the acidification line (14% decrease; 0.0127 ± 0.0111) followed by ambient (10% decrease; 0.0133 ± 0.0111) and warming (10% decrease; 0.0133 ± 0.0110) and finally OWA (7% decrease; 0.0138 ± 0.0112). These drops in diversity were similar between treatments, with the exception of the acidification line, which decreased more than other treatments (P < 0.001). The relatively small drop in diversity for OWA and warming lines was likely due to the higher minor allele frequencies of adaptive alleles; candidate SNPs were at higher minor allele frequency than the genome-wide distribution (Fig. S7). Thus, shifts in frequency of these more common variants could occur without purging variation from the populations.

## Discussion

The broad distribution, large population size, and variable coastal habitats inhabited by *A. tonsa* should, in theory, lead to the presence of adaptive genetic variation. Our results show that adaptation proceeded quickly under warming and OWA conditions and was dominated by selection (Fig. 2), indicating the presence of standing genetic variation for rapid adaptation. Further, the enrichment of candidate SNPs in coding regions of the genome indicates that protein coding variation, rather than regulatory variation, was the dominant target of selection and the main mechanism of adaptation across all of the selection regimes. This agrees with previous work focused on adaptation to acidification in urchins (Garrett et al., 2020) but is in contrast with other copepod experimental evolution that found links between allele frequency divergence and gene expression response (Griffiths et al., 2021). Others have found similar evolved shifts in gene expression during adaptation to global change conditions (Hsu et al., 2021; Mallard et al., 2020, 2018). Our previous work in this *A. tonsa* system has shown that gene expression rapidly evolved in response to OWA selection with strong signals of selection in regulators of gene expression such as transcription factors (Brennan et al., 2021). Thus, while the dominant signals of selection were found in coding regions, the evolution of regulatory elements influencing expression patterns are likely essential in the adaptation process.

In contrast to the rapid adaptation observed in OWA and warming conditions, the limited response to selection under acidification conditions suggests the presence of plasticity to tolerate pH stress. Previous work on *A. tonsa* has demonstrated that acidification alone can be tolerated through phenotypic plasticity (Langer et al., 2019) and that even OWA effects may be buffered by plasticity for a short period of time (Brennan et al., 2021). This plasticity is likely due to the high environmental variability in *A. tonsa* habitats, where even diel pH fluctuations occasionally exceed our acidification treatment (Baumann et al., 2015). Similarly, Vargas et al. (2017) recently demonstrated increased CO_2_-tolerance in copepods from high compared to low CO2-variability areas along the Chilean Pacific coast, suggesting that increased plasticity can evolve in variable habitats. Despite the likely presence of plasticity, we still observed a degree of selection and adaptation in the single stressor acidification conditions (Fig. 2). However, the contribution of drift to evolutionary change in this treatment was much larger than selection (Fig. 2). While this could be a true signal of drift, it is also possible that this is a signal of multiple, distinct selective pathways leading to adaptation, as has been observed in other systems (Barghi et al., 2019; Therkildsen et al., 2019). However, the targets of selection in acidification included functions commonly involved in resilience to pH stress (Fig. 6) and under selection in various species. For instance, we found enrichment for protein refolding, which is also under selection in mussels exposed to pH stress through development (Bitter et al., 2019). Similarly, cytoskeleton and actin regulation have been found to confer resilience in copepods (Bailey et al., 2017), purple sea urchin (Brennan et al., 2019) and oysters (Goncalves et al., 2017) exposed to low pH and these functions likely help stabilize cellular function in response to cellular stress.

Among copepod species, divergence in metabolic function and protein stabilization are common mechanisms promoting adaptation to higher temperatures (Kelly et al., 2017; Schoville et al., 2012). As we would predict, lines adapting to warmer conditions used mechanisms related to metabolic function and stabilization of proteins and cellular processes. Heat shock proteins are commonly expressed in response to heat stress to enable thermal resilience (Feder and Hofmann, 1999) and variation in their expression and protein coding sequence can confer increased heat tolerance (Sørensen, 2010; Tangwancharoen et al., 2018). In OWA and warming conditions there was a selected region in the gene *DNAJA3*, DnaJ heat shock protein family (Hsp40) member A3. *DNAJA3* regulates the expression of HSP70 (Faust et al., 2020), an inducible HSP that is responsive to temperature and general stress conditions (Mayer and Bukau, 2005). Selection for *DNAJA3* may be altering the sensitivity of HSP70 to environmental conditions. We also observed signals of selection for mitochondrial function conferring tolerance to high temperature. We find evidence of strong selection in two genes, NADH dehydrogenase 1 alpha and beta, that are mitochondrial and fundamental to the electron transport chain (Fig. S9). Variation in mitochondrial function plays a role in thermal tolerance as high temperatures can inhibit ATP synthesis (Chung and Schulte, 2020), as has been demonstrated in copepods previously (Harada et al., 2019). Our results suggest that adaptation to high temperature in *A. tonsa* may be countering energy limitation, though additional work would be needed to verify this hypothesis. Finally, heat stress can produce reactive oxygen species (ROS), which can lead to DNA, protein, and mitochondrial damage (Ali et al., 2006; Belhadj Slimen et al., 2014). In warming and OWA conditions we observed evidence of adaptation in focal adhesion kinase 1 (Fig. S9), a protein which can be activated by ROS and can influence downstream processes such as apoptosis (Ali et al., 2006; Ben Mahdi et al., 2000). While the functional consequences of this gene are unclear, it may maintain cellular function under the increased presence of ROS due to heat stress.

Processes facilitating adaptation to OWA conditions overlapped with warming and acidification, but were also unique, supporting the notion of a synergistic response to both stressors (Fig. 3). Responses to acidification and warming are metabolically costly, and the enrichment observed for metabolic processes in OWA conditions is expected. In acidification conditions this cost is driven by increased demands for acid-base balance, which requires the establishment of electrochemical gradients via ATP ion transporters to uptake bicarbonate from the environment (Whiteley, 2011). Previous work has shown that adaptation to acidification in copepods and other marine invertebrates includes functions related to ion transport (De Wit et al., 2016; Pespeni et al., 2013) while variation in acidification tolerance likely is linked to ionoregulatory capabilities (Lewis et al., 2013; Melzner et al., 2009). *A. tonsa* is euryhaline and can tolerate salinities ranging from hypersaline waters to freshwater (Calliari et al., 2008; Cervetto et al., 1999; Chinnery and Williams, 2004). Despite the predicted presence of plasticity for acidification tolerance, we found evidence for adaptation in OWA lines, and to a lesser extent under acidification alone. Of particular interest was the strongly selected gene Na^+^,K^+^-ATPase subunit α-4 (*NKA* α4), a gene underlying salinity adaptation in the copepod *Eurytemora affinis* (Fig. S9) (Gerber et al., 2016; Stern and Lee, 2020) as well as killifish (Brennan et al., 2018). Here, allele frequency change in *NKA*, is likely related to improvements of acid-base regulation and/or energetic efficiency in warm, acidified environments.

The strong signals of selection at F25 were also present after just three generations of selection in OWA conditions where adaptive alleles were more likely to shift towards the adaptive state by F3 than neutral loci (Fig. 5). Our previous results showed that, following a decrease in fitness at F0, OWA animals largely recover fitness by F3 (Dam et al., 2021). While some of this response may be due to transgenerational plasticity, the allele frequency shifts at F3 suggest that adaptation played a role in this recovery. Over the course of just a single or few generations, adaptation can proceed in both wild (Bumpus, 1899; Campbell-Staton et al., 2017; Donihue et al., 2018) and lab populations (Bitter et al., 2019; Brennan et al., 2019; Garrett et al., 2020; Pespeni et al., 2013). Our results support these findings and show that short term adaptive changes can occur over only a few generations and can be indicative of longer-term evolutionary responses. Thus, even short-term generation selection experiments can be informative about the adaptive potential and responses across multiple generations.

Under a hard sweep scenario, we would expect pronounced reductions in genetic diversity that would likely scale with the strength of selection (Pennings and Hermisson, 2006). However, the lack of connection between the strength and response to selection with genetic diversity coupled with the intermediate frequencies of selected alleles indicates that selection proceeded via soft, polygenic sweeps (Figs. S6, S7). From a quantitative genetic perspective, polygenic selection is likely to result in relatively small allele frequency shifts at many loci (Pritchard et al., 2010), a finding that our data support with the broad allele frequency responses across the genome coupled with few loci with large shifts (Fig. S2). Simultaneously, variants at higher frequency are more likely to fix during adaptation and thus may play an important role during the adaptation process (Hermisson and Pennings, 2005; Messer et al., 2016). These intermediate frequency loci are likely maintained by balancing selection (Höllinger et al., 2019) perhaps due to naturally fluctuating environmental conditions in typical coastal habitats inhabited by *A. tonsa* (Baumann and Smith, 2018; Nixon et al., 2004). Indeed, it appears that rapid adaptation commonly targets alleles at intermediate frequency (Barghi et al., 2019; Brennan et al., 2019; Kelly and Hughes, 2019; Mallard et al., 2018). In particular, Stern and Lee (2020) found that balancing selection maintained standing genetic variation that was selected upon to enable the invasion of and rapid adaptation to freshwater environments in the estuarine copepod *Eurytemora affinis*. It is likely that a similar balancing selection mechanism is involved in maintaining the adaptive genetic variation we observe here.

The application of covariance-based analyses revealed selection was driving the allele frequency changes across the different selective regimes and these methods represent an advance in our ability to disentangle drift from selection in temporal datasets that are adaptive via polygenic selection (Buffalo and Coop, 2020, 2019). However, there are limitations with our analyses that are important to consider. First, both covariance and frequency change methods, such as CMH, identify targets of selection that are consistent across the replicates in each treatment. This assumption does not always hold as polygenic traits can be genetically redundant and adaptation can proceed through unique pathways (Barghi et al., 2020; Láruson et al., 2020). As a result, our identification of candidates and the proportion of allele frequency change due to selection are prone to false negatives; there are likely adaptive loci that are unique targets of selection in only a subset of replicates. Conversely, linked selection and LD can lead to false positives in the candidate gene approach, inflating the number of candidates (Tobler et al., 2014).

Controlling for lab adaptation in experimental evolution and E&R is essential when inferring shared responses to selection. When lab effects were not removed, we falsely increased the overlap in candidate SNPs (Figs S2) and the shared response to selection between treatments (Fig. 3). Our extension of Buffalo and Coop’s covariance methods (Buffalo and Coop, 2020) to account for lab adaptation signals revealed that all of the shared variance between acidification and warming as well as the majority between acidification and OWA was due a shared response to lab conditions, lab adaptation (Fig. 3). Despite the widespread acknowledgement of the artificial nature of lab environments, which is sure to exert a selective pressure, surprisingly few studies include an appropriate control to account for these effects. Our demonstration of the removal of the signals of lab adaptation, through both the covariance and candidate SNP analyses, should serve as a starting point for others to account for these effects in future work.

The results from our full-factorial evolution experiment demonstrate the complexity of predicting responses to global change conditions when multiple environmental parameters are simultaneously shifting. We show that acidification alone had a limited impact on fitness and that selection weakly affected allele frequencies across the genome. However, when coupled with warming to create OWA conditions, the response to selection was non-additive and synergistic; only 1% and 37% of the variation in allele frequency changes in OWA lines was shared with acidification or warming, respectively (Fig. 3). Similarly, of the response to selection in OWA conditions, 47% was not shared with either the acidification or warming selection regimes. These results suggest that a large proportion of the OWA response to selection is unique from either individual component of the selection regime. This has consequences for our ability to predict targets of selection and resilience from genomic data in natural populations. Even relatively benign stressors, such as change in pH for copepods, can have important impacts when paired with other abiotic changes. As such, predicting resilience or adaptive potential becomes more difficult as the number of changes to the environment increases. Conversely, the shared response to selection and common functional targets between OWA and the single stressors suggest that functionally, many of the responses are related. Finally, the large shared signal of selection between OWA and warming suggests that adaptive potential for increased temperature will be a dominant component enabling resilience as the climate continues to change.

## Methods

### Selection experiment

The details of the selection experiment have been reported in detail previously in Dam et al. (Dam et al., 2021). This experiment was conducted with four replicates per treatment designed to simulate future ocean conditions that were warmer and more acidic with the following target environments: 1. Ambient (18°C, 400 μatm CO_2_, pH ^~^8.2); 2. Acidification (18°C, 2000 μatm CO_2_, pH ^~^7.5); 3. Warming (22°C, 400 μatm CO_2_, pH ^~^8.2); 4. OWA (22°C, 2000 μatm CO_2_). These levels were chosen as ambient conditions represent current conditions in the native habitat while elevated CO_2_ and temperature align with predicted future conditions in the year 2100-2300 (Bindoff et al., 2019; Caldeira and Wickett, 2003). Four replicates is generally considered suitable to achieve good power in E&R experiments (Kofler and Schlötterer, 2014). Temperature treatments were achieved using four incubators, two for 22°C, the other two for 18°C, while elevated CO_2_ was controlled by continuously mixing CO2 directly to the bottom of each replicate. Ambient CO_2_ conditions were achieved by mixing CO2-stripped air into each culture. Dissolved oxygen was > 8mg L^−1^ for all cultures.

Adult *Acartia tonsa* (n=1,000) were collected in June of 2016 from Esker Point Beach, Groton, CT, USA (41.320725°N, 72.001643°W). After at least three generations of lab acclimation, eight replicate cultures were each started with 160 females and 80 males. Four of these eight cultures had temperature increased by 1 degree Celsius per day while the others remained at ambient temperature. After temperature acclimation, adults were allowed to lay eggs, producing an average of 7,173 eggs per replicate. The warm treatments replicates (four of each for OWA and warming) were seeded by the warm acclimation animals (average of 3586/replicate) and the ambient temperature treatments by the ambient animals. Cultures were fed equal concentrations of phytoplankters *Tetraselmis* sp., *Rhodomonas* sp., and *Thalassiosira weissflogii* every 2-3 days at food replete condition (≥800 μg Carbon L^−1^), following established protocols (Feinberg and Dam, 1998). Cultures were reared for 25, non-overlapping, generations. Samples for genomic analysis were collected from each replicate from the ambient treatment at F0 to represent the starting population allele frequencies and from each of the selection regimes, including ambient, at F25. At generation F3 samples were also collected from the OWA treatment. However, due to an experimental mishap the F3 replicates were inadvertently mixed. While this removes the ability to estimate replicate specific responses, we could still accurately estimate mean F3 allele frequencies.

### Genome annotation

We based genomic analysis on the publicly available, yet unannotated, *Acartia tonsa* genome (Jørgensen et al., 2019). We followed the GAWN annotation pipeline (https://github.com/enormandeau/gawn) to complete the annotation of the existing assembly. GAWN takes an evidence-based approach to assembly, using an assembled transcriptome as the evidence for gene models; we leverage the high quality transcriptome from Jørgensen et al. (2019) for this purpose. Briefly, this method uses GMAP (Wu and Watanabe, 2005) to align the transcriptome to the genome, transdecoder (Haas et al., 2013) to determine open reading frames, and blastx (Camacho et al., 2009) to annotate predicted genes.

### SNP identification

Capture probes were designed to target regulatory and coding regions across the genome. Regulatory probes were located within 1000 bp upstream of transcription start sites and coding probes were located in exons; both regulatory and exonic probes were chosen to maximize capture quality. This process resulted in 32,413 probes (21,311 exonic, 11,102 regulatory). DNA library preparation and sequencing was conducted by Rapid Genomics (Gainesvill, FL, USA) on a HiSeq 4000 with 150 bp paired end reads.

Raw reads were trimmed for quality and adapter contamination with Trimmomatic v0.36 (Bolger et al., 2014) and mapped to the *A. tonsa* reference genome (Jørgensen et al., 2019) with BWA-MEM (Li, 2013). Raw genomic data is deposited in GenBank: BioProject number PRJNA590963. Variants were called using Varscan2 (Koboldt et al., 2012) using liberal thresholds of minimum variant frequency of 0.01, p-value threshold of 0.1, minimum alternate reads 2, and minimum coverage of 30x, resulting in 10,368,816 sites. Following variant calling, sites were stringently filtered to include only those where coverage was > 40x in all samples, minor allele frequency > 0.01 in at least 4 samples (i.e., one treatment), including only biallelic sites, and excluding sites with depth per sample above the 99% quantile (912x) to control for mismapping. This strict filtering resulted in a final set of 394,667 variant sites.

### Characterizing genetic variation

We summarized genome-wide variation in allele frequencies between all samples using principal component analysis with the prcomp function in R. To understand the demographic impacts of the selection regimes, we estimated levels of linkage disequilibrium (LD) and genetic diversity (π) for all replicates. LD was estimated using LDx, a method developed for pool-seq data that leverages haplotype information from single reads to estimate linkage between pairs of SNPs over short distances (Feder et al., 2012). We estimate the decay of LD by regressing the log of physical distance with LD between base pairs; to estimate the slope and intercept of each treatment we include replicate as a random effect with the R package lme (Bates et al., 2014).

Genetic diversity was estimated using Popoolation^59^. We identified 1,940 100 bp windows in 987 unique scaffolds across the genome that were present across all samples. We estimated π in 100bp sliding windows with a 100bp step size. Each window required a minimum coverage of 15x, max coverage of 1000× (to avoid mapping errors), and at least 0.5 of the window meeting these thresholds. Resulting windows were required to be sequenced across all samples. To take into account the independent replicates within each treatment, we used pairwise Wilcoxon Rank Sum tests with a Holm correction for multiple testing. All statistics were performed in R^49^.

### Clade assignment

*Acartia tonsa*, while broadly distributed, is made up of mitochondrial lineages that likely represent cryptic species (Chen and Hare, 2011, 2008; Figueroa et al., 2020), at least two of which are reproductively isolated (Plough et al., 2018). Therefore, it is important to understand the mitochondrial lineage of the populations used here and if selection for different lineages could be driving the responses to selection we observe. We reconstructed COI haplotypes present in the starting and evolved cultures to ensure different signals of adaptation between conditions were not simply shifts in the cryptic species composition of the cultures across time. We obtained the reference COI sequence from Figueroa et al. (2020) and aligned our raw fastq files to this reference, calling variants in the same manner as the full dataset, above. The distribution of allele frequencies of these variants was bimodal, with nearly all variants close to fixation (Fig. S10). We generated a consensus sequence for the major and minor variant for each sample, generated phylogenetic trees using Mr. Bayes (Ronquist et al., 2012), including samples from Figueroa et al. (2020) and following their analysis. We then determined the mitochondrial lineage by comparing the clade in which our samples fell to Figueroa et al. which found that these copepods used here were clade X from Figueroa et al. (Fig. S11)(2020). These results were validated by sequencing COI from individual copepods following Figueroa et al. from these same cultures at approximately generation 100. At F0 samples were dominated by haplotypes from a single clade with a very low frequency of haplotypes from a sister clade (Fig. S10). By F25, the minor haplotype was largely absent across all samples. Thus, any differential signals of adaptation between experimental conditions are not due to the presence of cryptic species in the samples but rather selecting with each condition on the same lineage’s genetic variation. See the supplemental methods for more details.

### Estimates of allele frequency change within and between treatments

Disentangling the extent to which the variance in allele frequency change within and between treatments is due to drift or selection is difficult due to the subtle changes that can occur during polygenic adaptation. To overcome this limitation, we used *cvtk* (https://github.com/vsbuffalo/cvtkpy) (Buffalo and Coop, 2020, 2019). This approach uses the replication within treatments to calculate the covariance between samples and partition the variance in allele frequency change from F0 to F25 into selection and drift components. That is, consistent temporal allele frequency changes within a selection regime are due to selection and inconsistent shifts are drift. Further, this framework can be used to assess the convergent correlation between different treatments to quantify the degree to which selection targets common genomic regions in different conditions, generally revealing the consistency of allele frequency changes between each treatment. Finally, we extended *cvtk* to incorporate signals of lab adaptation where the ambient control should only be selected for lab conditions. Because of this, any covariance between this control and our selection lines indicated adaptation to the lab environment and not a particular selective pressure. We calculated the covariance of allele frequency change from F0 to F25 in 10,000 basepair windows for each replicate and used these covariance estimates to calculate the proportion of the change of allele frequency due to drift, selection, and lab adaptation; results hold when window size is changed to 1,000 basepairs. Uncertainty of these estimates was determined with bootstrap resampling of windows. See the supplemental materials for a detailed discussion of these methods.

We determined specific loci evolving due to selection by simulating the expected drift over 25 generations using the poolSeq packing in R (Taus et al., 2017) and following Barghi et al.(2020). Using the starting allele frequencies at F0, we mirror our experimental design simulated allele frequency trajectoires for four replicates across 25 generations with no selection under a Wright-Fisher model. We first estimated the mean effective population size to be 414 across all replicates. We then simulate allele frequency changes using this effective population size, a census size of 3,000, sample size of 50 individuals, and a simulated sequencing depth of 167 at F0 and and 106 at F25, matching our observed depth. This simulation was repeated 500 times and for each replicate we used Cochran–Mantel– Haenszel (CMH) tests in the Popoolation2 package (Kofler et al., 2011) to generate a null distribution of neutral allele frequency change. We calculated empirical p-values for our observed data from this simulated distribution using empPvals in the q-value R package (Storey et al., 2019). We identified candidate loci underlying adaptation as those with empirical p-values less than 0.05. Significance of overlap between sets of candidate SNPs between treatments was calculated using SuperExactTest in R (Wang et al., 2015).

Functional enrichment was tested using TopGO v. 2.36.0 (Alexa et al., 2006; Alexa and Rahnenfuhrer, 2019) with the weight01 method with terms that had at least 5 annotated genes. We used GOSemSim (Yu et al., 2010) with Wang’s method in R to calculate the similarity between terms with p-values < 0.05 and converted this similarity matrix to a dissimilarity matrix for hierarchical clustering and plotting using ggdendro in R (de Vries and Ripley, 2016).

## Acknowledgements

Funding: This work was funded by the National Science Foundation grants to M.H.P. (OCE 1559075; IOS 1943316) and H.G.D, H.B. and M.F. (OCE 1559180) as well as Connecticut Sea Grant (R/LR-25) awarded to H.G.D., M.F. and H.B. We thank Jon Puritz for helpful feedback on this manuscript.

## Data availability

Sequence data are available on NCBI BioProject number PRJNA590963. Data files and supplemental tables can be found on Zenodo: https://doi.org/10.5281/zenodo.5093797. Code to run all analyses can be found on github: https://github.com/rsbrennan/tonsa_25_gen_ER

## Competing Interests

The authors declare no competing interests.

## Supplemental Materials

### Mitochondrial lineage assignment

To ensure the selection experiment did not confound selection between mitochondrial lineages, which are potentially cryptic species, and selection within a lineage, we determined the major and minor mitochondrial haplotypes present in our samples. This analysis follows previous COI work conducted in *A. tonsa* (Chen and Hare, 2011, 2008), particularly Figueroa et al. (2020). We used a reference COI from FIgueroa et al. to align and pull out the COI region in our raw FASTQ files. Variants were called using Varscan2 (Koboldt et al., 2012) in the same manner as our full dataset. Because these are pooled data, we cannot determine haplotypes of individuals. Further, if two lineages were present at equal frequencies, it would be difficult to determine which two haplotypes were present; the variants of each would be mixed in these pooled data. We assessed the distribution of allele frequencies to predict the number and frequency of mitochondrial haplotypes present in these data. Given the bimodal distribution of variants, where SNPs are either nearly fixed or absent (Fig. S10), there are likely two mitochondrial lineages present in these samples; one at very high frequency, and the other at low frequency. Looking at the shift in variant frequency from F0 to F25, it is apparent that the low frequency variant drops out over time while the high frequency variant (which is nearly fixed) remains. Because of the low number of variant sites, the high frequency variant also appears to match the reference sequence, which is from an individual from clade X. All results below hold when a reference from a different clade is used instead.

We reconstruct the major and minor haplotypes present in our data. Given the bimodal distribution of variants, we can reconstruct the consensus sequence of the major haplotypes as variants that are nearly fixed, while the minor haplotype consist of the low frequency variants. We only construct the minor haplotypes for F0 samples, as these haplotypes are no longer present by F25. Variants were filtered in R and VCF files containing either the major or minor variants were created. *bcftools* was used to generate consensus sequences for each sample and haplotype and sequences were concatenated with fasta sequences from Figueroa et al. to allow for direct comparisons downstream. This fasta file was aligned with MUSCLE (Edgar, 2004), converted to NEXUS format in R using APE (Paradis et al., 2004), and a phylogenetic tree was built using MrBayes (Ronquist et al., 2012) using the substitution model HKY + I + G, following Figueroa et al. The plot was generated with ggtree in R (Yu et al., 2017). The resulting tree clearly shows that the major haplotype is clade X, with a low frequency of clade S present at F0 (Fig. S11). The S clade drops out by F3 and is not present in any samples at F25, including the ambient selection line. Given the low frequency of clade S and its absence from all lines by F25, the divergence detected in our experiment is not simply due to selection for a mitochondrial clade, but due to selection within individuals of clade X.

### Analysis of covariance in allele frequency change

The replicated nature of our experiment coupled with the multiple selection regimes allows us to disentangle the relative contributions of drift and selection on the genome-wide changes in allele frequencies. We use and expand upon the approach developed by Buffalo and Coop (Buffalo and Coop, 2020). This method quantifies genome-wide covariance in the change in allele frequencies between replicates of a single treatment and between treatments to determine the relative contributions of drift and selection as well as to assess the degree of shared selection between selection regimes. Finally, we can leverage the presence of the control ambient line to estimate and remove the effects of the aforementioned estimates of lab adaptation.

First, it is possible to partition the changes in allele frequencies within a treatment into selection and drift components for replicate A as:

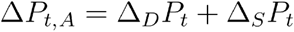

where Δ_*S*_*P*_*t*_ is the change due to selection and Δ_*D*_*P*_*t*_ is drift. The proportion of change due to selection can be further defined as the allele frequency change due to lab selection common to all replicates within a treatment group, Δ_*L*_*P*_*t*_, as well as the change due to selection within a specific selection regime, Δ_*R*_*P*_*t*_. The change in allele frequency in replicate A can be partitioned into,

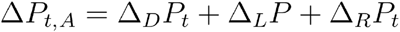

Because the terms above are uncorrelated, the variance is,

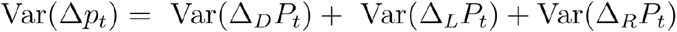

We estimate the shared effects of selection regime from the allele frequency changes as the covariance of allele frequency change between any two replicates,

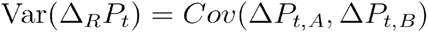

Where A and B indicate different replicates within a selection regime. Thus, to estimate the total shared response to selection within a selection regime, we estimate the covariance between all pairwise replicates and take their mean. Further, because of the presence of an ambient control line in our study, we can estimate the contribution of adaptation to the lab environment to the overall changes in allele frequency. The shared variance between a selection regime and the control represents the contribution of lab adaptation within a treatment group, again estimated as the mean of all pairwise comparisons between the two groups, giving VarΔ_*L*_*P*_*t*_. We can subtract this value from the shared response within a selection regime, Var(Δ_*R*_*P*_*t*_), to get an estimate of the response to selection that is independent of estimated average lab selection effect.

After determining the contributions from selective regime and lab adaptation, the remaining variance can be attributed to the drift component. Finally, these values can be divided by the total variance to find the proportion each contributes to the overall variance in change in allele frequency.

Next, we can use similar principles to determine the shared response to selection between each selective regime. Here, the shared response to selection is again the covariance in allele frequency change between any two replicates, as above, but now from two different treatments,

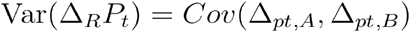

We take the mean of all possible pairwise comparisons between treatments and scale this by the total variance to determine the proportion of total variance of allele frequency change that is shared. Finally, lab adaptation can be estimated and accounted for as described above.

## Accounting for shared variance due to limited F0 replication

In these data, covariances are calculated from the change in allele frequency from the same set of F0 samples for all treatments. While these F0 samples likely represent the pre-selective state of the population, replicating these samples in this way leads to a spurious increase in the covariance estimates between samples in different treatments due to shared sampling variance. This is made clear in the covariance heatmap (Fig. S1A) where along the diagonal where the same replicate is compared between treatments the covariance is increased; Fig. S1B shows these covariance calculations in a different format where the same pattern is evident. Similarly, we see the same pattern when the convergence correlation is calculated (Fig. S12). To avoid this inflation, we dropped any covariances between samples with the same F0 reference when calculating shared response to selection. For example, the covariance in allele frequency change between OWA replicate 1 and Acidification replicate 1 would not be included when quantifying the shared selection response between these treatments. While this reduces our replication to an extent, the estimates are likely much more accurate estimates of the true impact of selection on allele frequency changes as all covariances are independent as a result.

**Supplemental table 1:** Number of loci assigned to each functional category for each SNP set. The first value is the total number of loci, the 2nd is the proportion of the total. P-values from chi-squared tests relative to the genome-wide distribution. Underlined values are significant (Bonferroni correction: P < 0.05/15 = 0.0033). The “No Annotation” category refers to loci that were not on a scaffold with any annotated genes, indicating they are likely in fragmented regions of the genome assembly.

|  | All loci | OWA candidates | Warming candidates | Acidification candidates |
| --- | --- | --- | --- | --- |
| **No annotation** | 54,228<br>0.137 | <u>1,384</u><br><u>0.072 (p << 0.001)</u> | <u>908</u><br><u>0.068 (p << 0.001)</u> | <u>339</u><br><u>0.0997 (p << 0.001)</u> |
| **Downstream** | 28,197<br>0.071 | <u>1,207</u><br><u>0.063 (p << 0.001)</u> | 925<br>0.070 (p = 0.42) | 249<br>0.073 (p = 0.70) |
| **Exon** | 229,898<br>0.583 | <u>12,691</u><br><u>0.665 (p << 0.001)</u> | <u>8922</u><br><u>0.670 (p << 0.001)</u> | 2105<br>0.619 (p = 0.029) |
| **Intron** | 46,837<br>0.117 | 2,188<br>0.115 (p = 0.13) | <u>1437</u><br><u>0.108 (p = 0.001)</u> | 396<br>0.116 (p = 0.73) |
| **Promoter** | 35,507<br>0.0899 | 1627<br>0.085 (p = 0.039) | 1117<br>0.084 (p = 0.028) | 311<br>0.091 (p = 0.78) |

## Supplemental Figures

**Figure S1:**
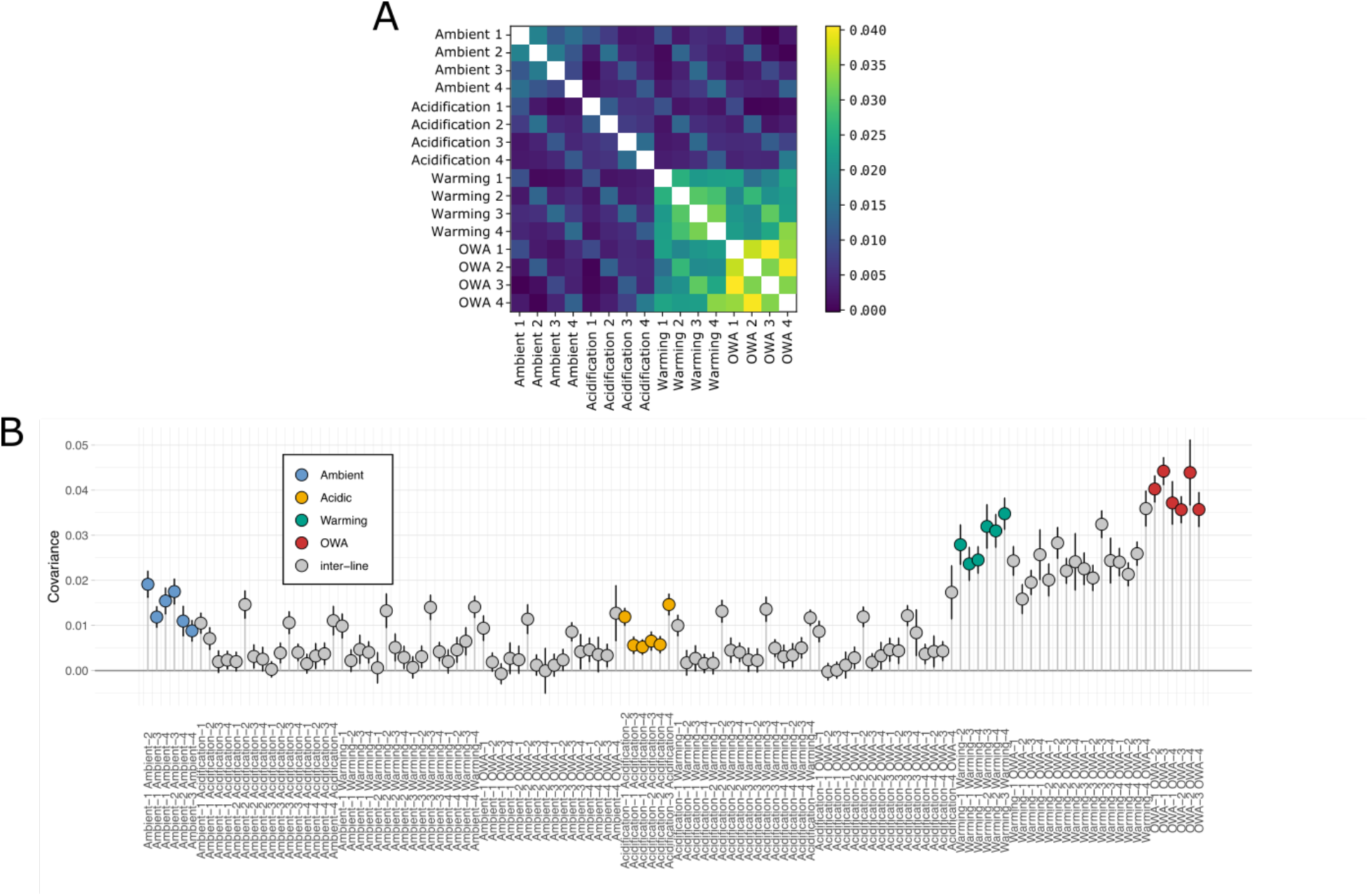
Pairwise covariance in allele frequency change between samples from F0 to F25. A) Each square and color indicate the covariance between two samples. Above the diagonal is a mirror of below. B) The same covariance estimates as in A, but with 95% bootstrap confidence intervals. Color indicates the group comparison. Note the increase in covariance between samples sharing the same replicate number due to an artefact from calculating allele frequency change from the same F0 sample. Samples with shared F0 samples (i.e., the same replicate number) were dropped from further calculations to avoid this bias.

**Figure S2:**
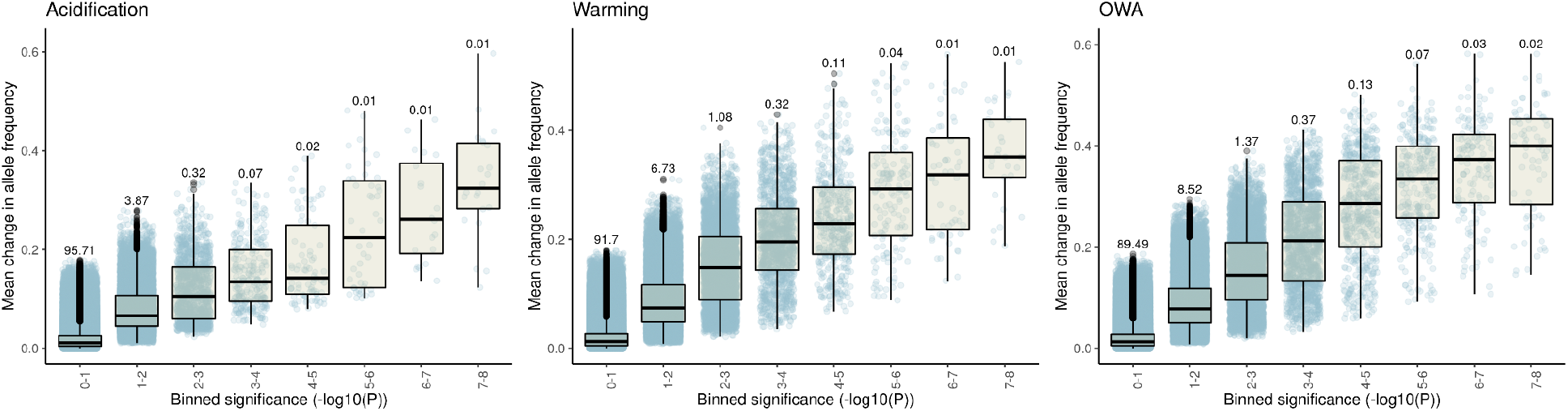
Empirical p-values from drift simulations. Points are individual loci with Tukey’s boxplots. Values above each boxplot indicate the proportion of total loci in that bin.

**Figure S3:**
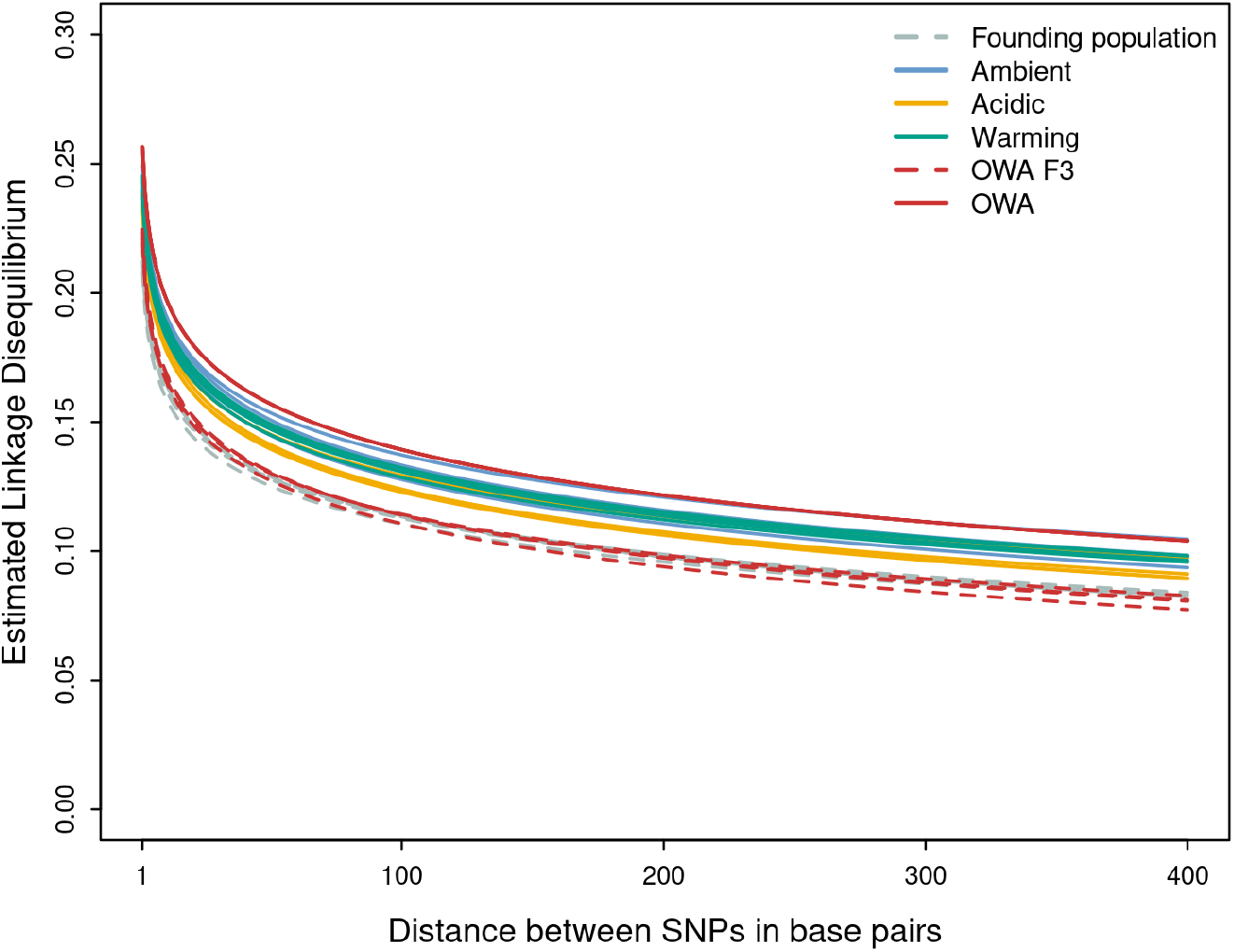
Linkage disequilibrium estimates. Decay curves were fit by regressing the log of the physical distance with LD between base pairs. LD estimates increased with the strength of selection relative to the F0 founding population (P < 0.001). Founding population: intercept = 0.212 ± 0.001, slope = −0.0497 ± 0.0003; Ambient: 0.245 ± 0.002, −0.0562 ± 0.0003; Acidification: 0.235 ± 0.001, −0.0549 ± 0.0003; Warming: 0.24181834 ± 0.001, −0.0556 ± 0.0003; OWA: 0.251 ± 0.002, −0.058 ± 0.0003; OWA F3: 0.222 ± 0.001, −0.0544 ± 0.0002

**Figure S4:**
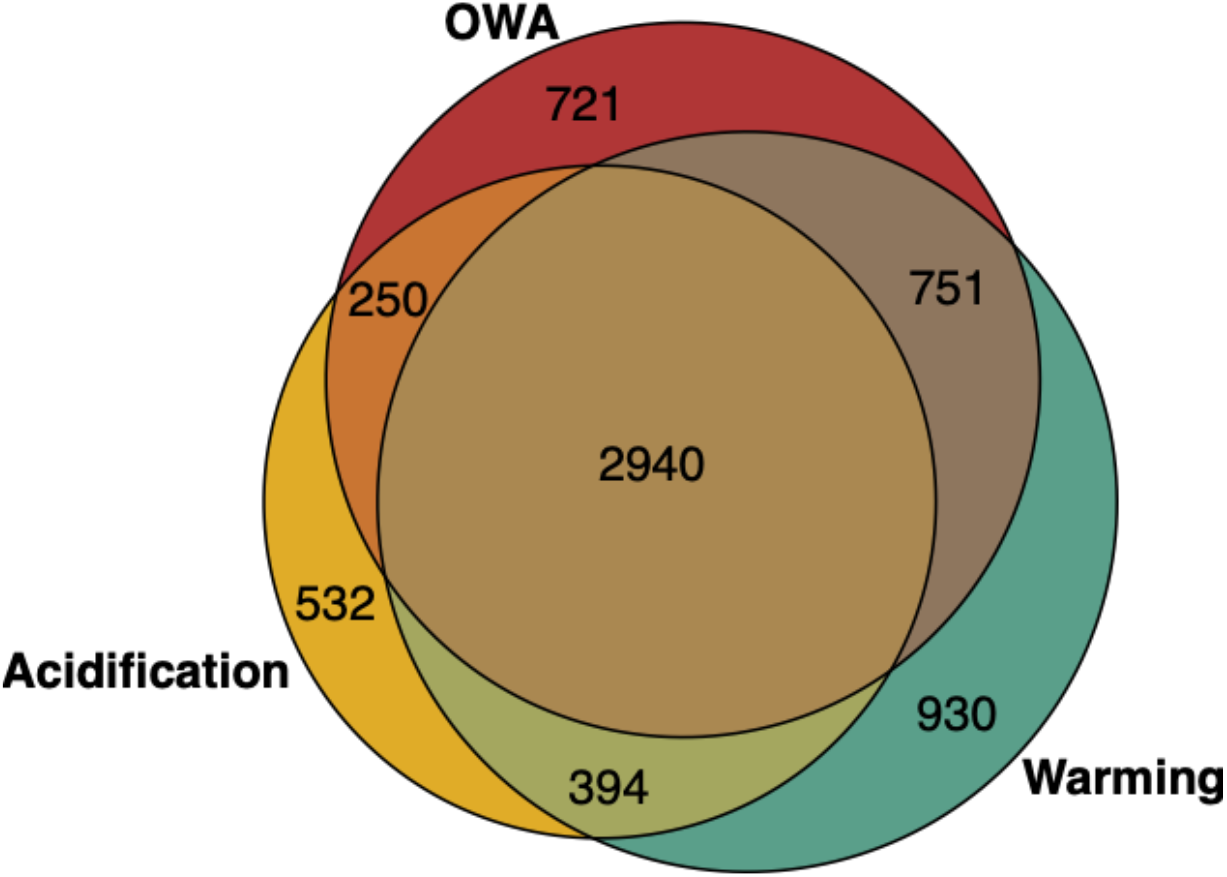
Venn of lab adaptation loci. Loci that are consistently changing allele frequency greater than expected under drift in ambient condition as well as any of the treatments. The core set of lab adaptation loci are highly overlapping between the three treatment groups.

**Figure. S5:**
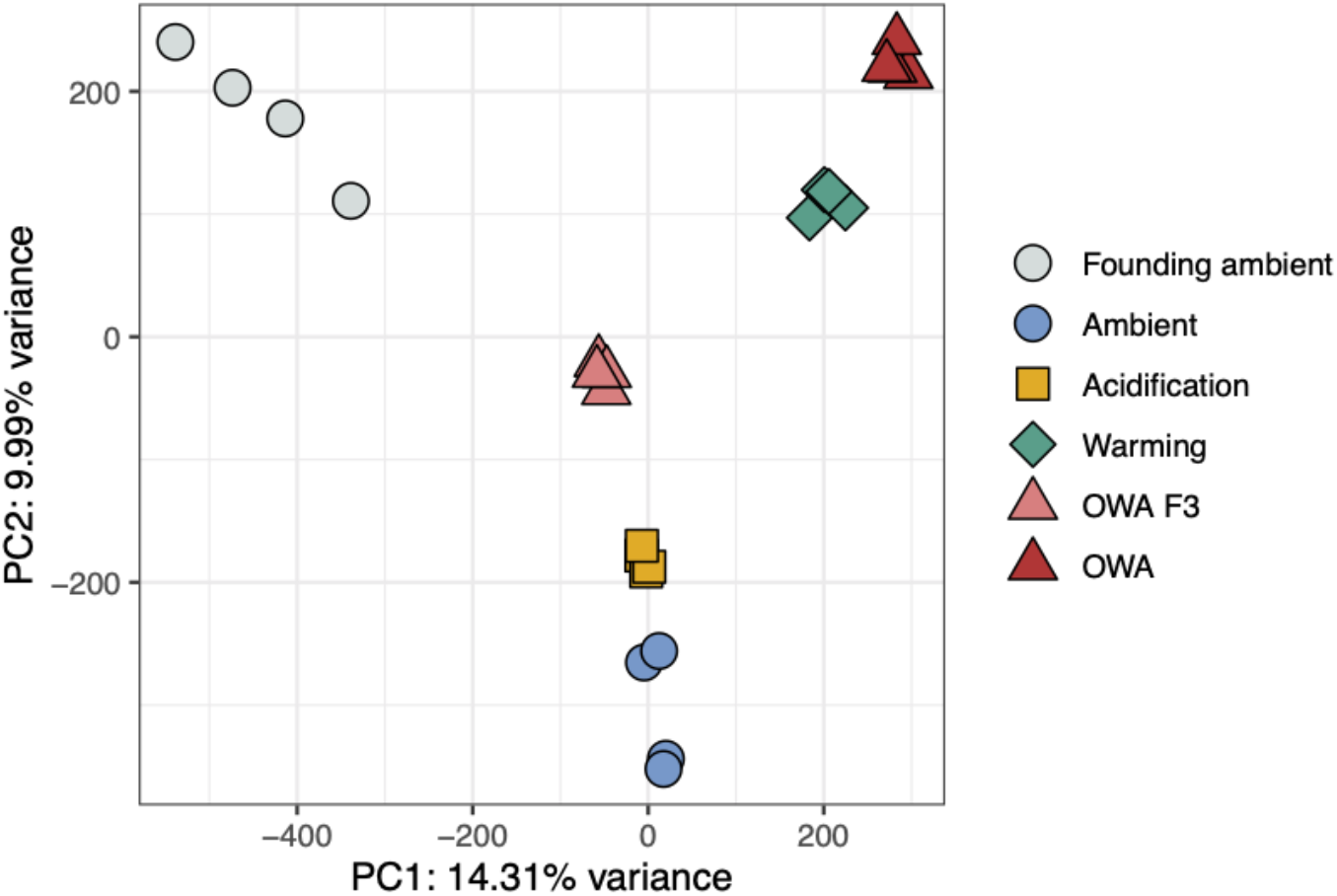
PCA with greenhouse F3 samples. Note that greenhouse samples at F3 are pseudo-replicates that were sequenced from the same pool of individuals that were inadvertently mixed from all four replicates.

**Fig. S6:**
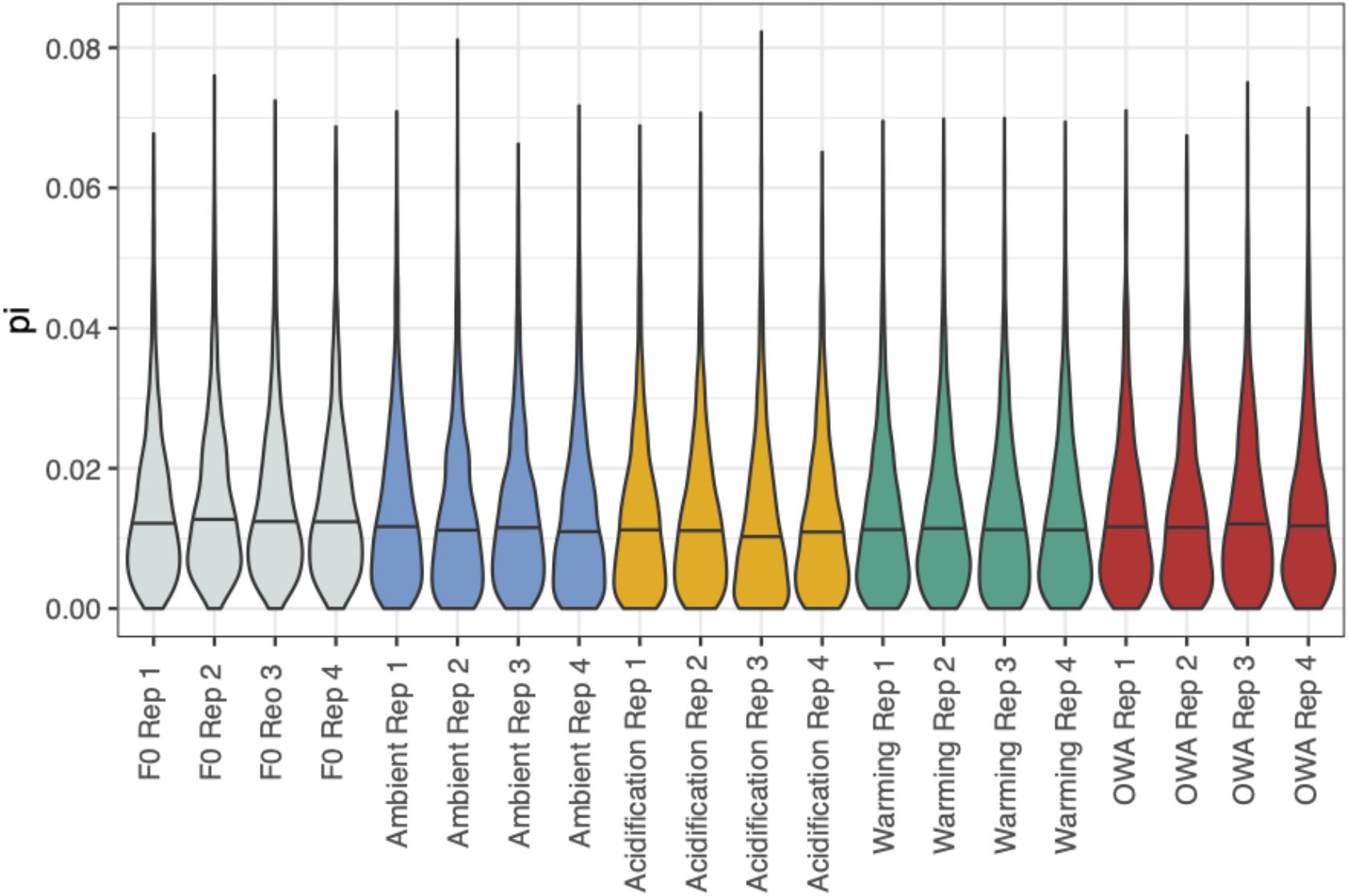
Genetic diversity estimates. All treatment groups lost genetic diversity relative to the founding F0 population (Wilcoxon signed-rank test, P < 0.0001). Estimates for each group: F0 founding population: 0.0148 ± 0.0111; ambient: 0.0133 ± 0.0111; acidification: 0.0127 ± 0.0111; warming: 0.0133 ± 0.0110; OWA: 0.0138 ± 0.0112. Between the F25 treatments, only acidification lines were significantly lower than other lines (P < 0.001).

**Fig. S7:**
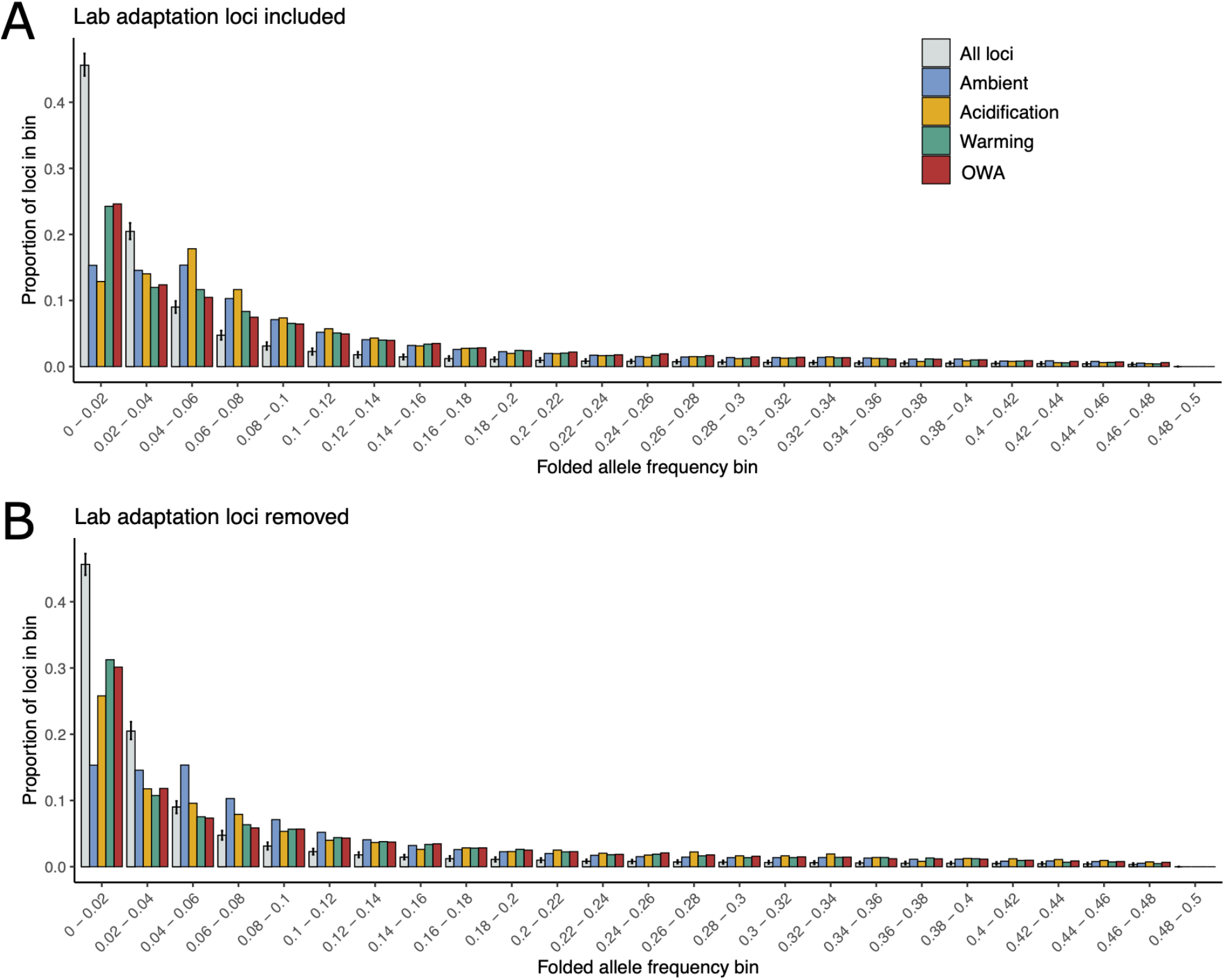
Starting minor allele frequencies at F0 for loci significant for different treatments. Panel A does includes lab adaptation loci in each group while these are removed in panel B. Error bars around the all loci group are 95% confidence intervals from 1000 random samples from all loci where the number of loci sampled was equal to the number of significant Acidification loci (A: 7,516, B: 3,400). Because the acidification group had the least number of significant loci, these confidence intervals are more broad than if sampling was based on warming or OWA values.

**Figure S8:**
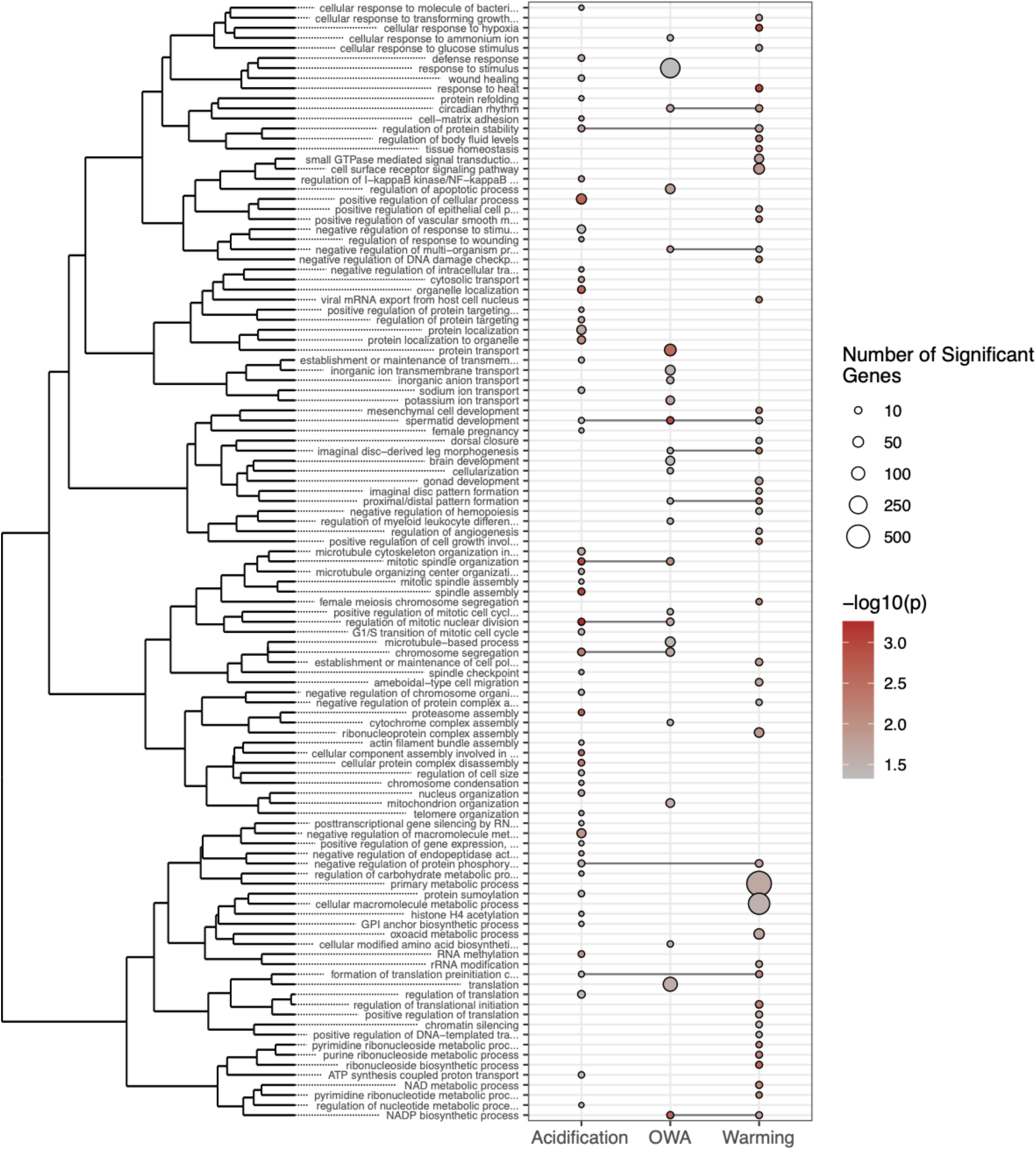
Gene ontology (GO) enrichment from candidate SNPs under selection. The size of a point indicates the number of genes in a category and color shows p-value. GO terms are clustered by similarity and the presence of a dot in a column indicates significant enrichment for that term in the treatment condition.

**Figure S9:**
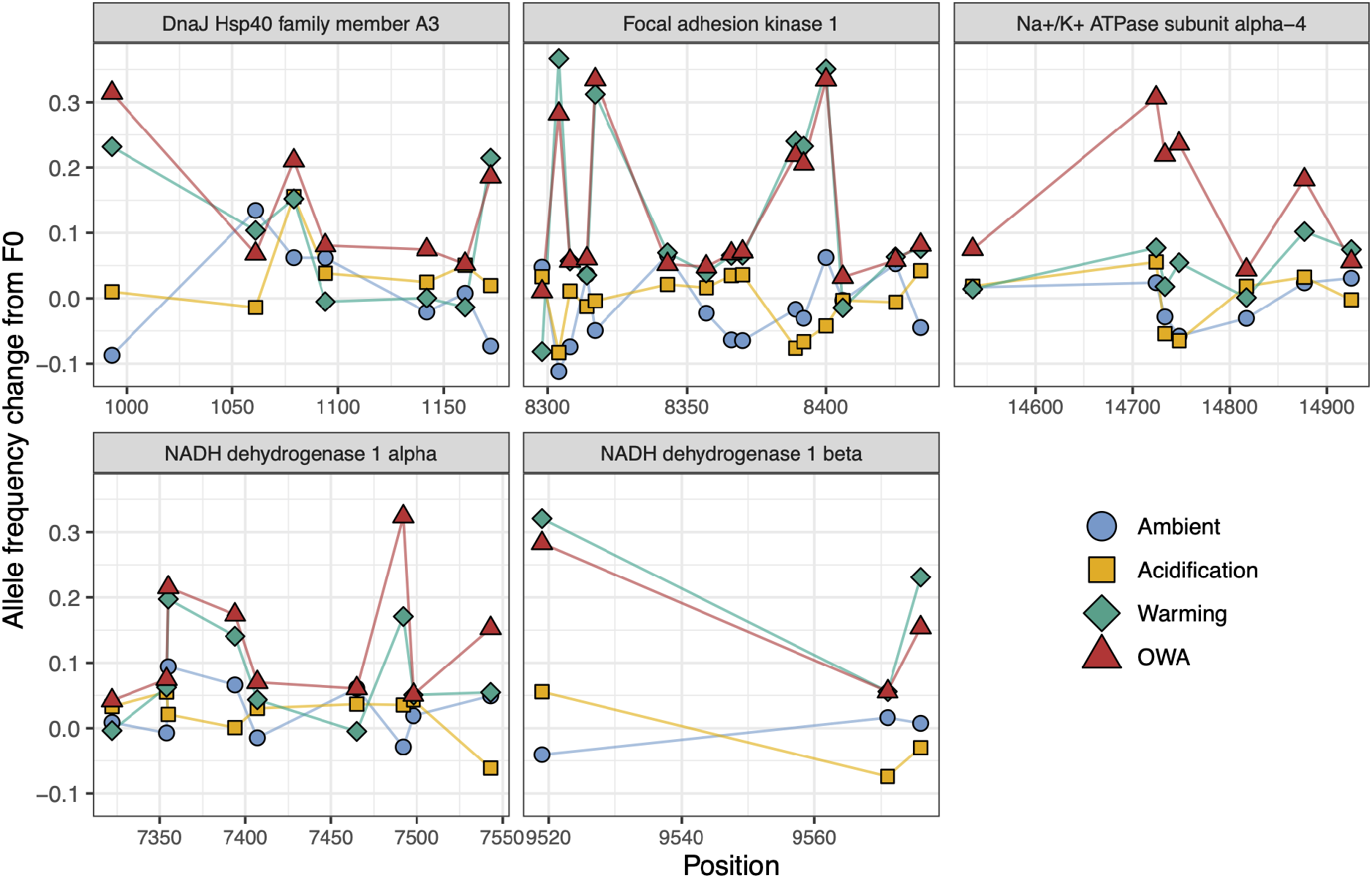
Representative candidate genes underlying rapid adaptation. Points represent the average allele frequency change among replicates from the starting frequencies at F0. Loci have been filtered for minor allele frequency > 0.05 at F0.

**Figure S10:**
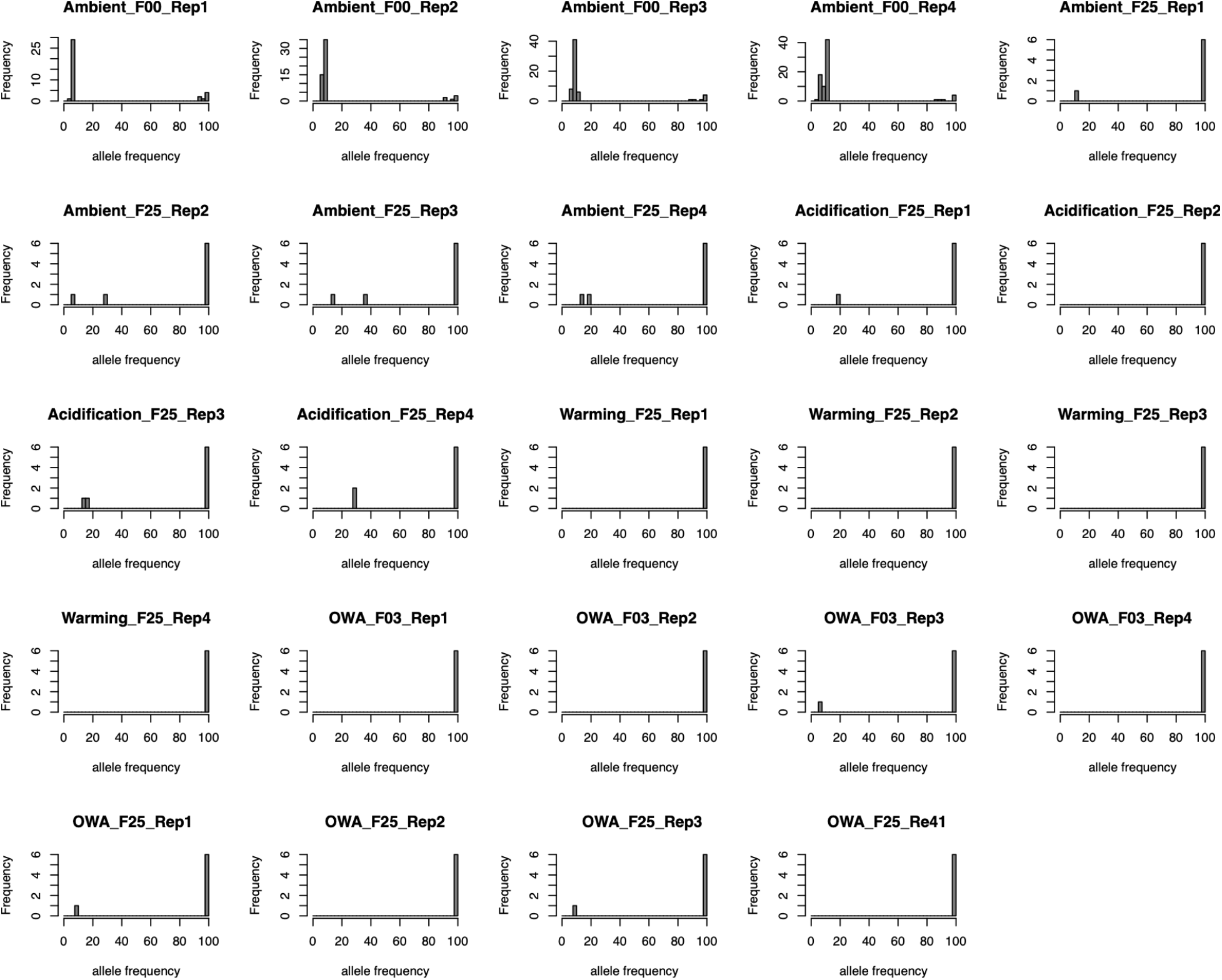
Frequency of mitochondrial haplotypes. Histograms show the frequency of haplotypes from variants in the pooled data. At F0, there were low frequency variants present in the data, indicating the presence of a low frequency haplotype. These minor alleles drop out by F3 and nearly absent by F25 across all samples. This indicates that the vast majority of samples matched the reference genome after 3 generations of selection. Given the low starting frequency of minor mitochondrial haplotypes, this suggests the pooled samples likely consisted of a dominant mitochondrial haplotype and selection across the the rest of the genome was likely not due to shifting frequencies of mitochondrial clades.

**Figure S11:**
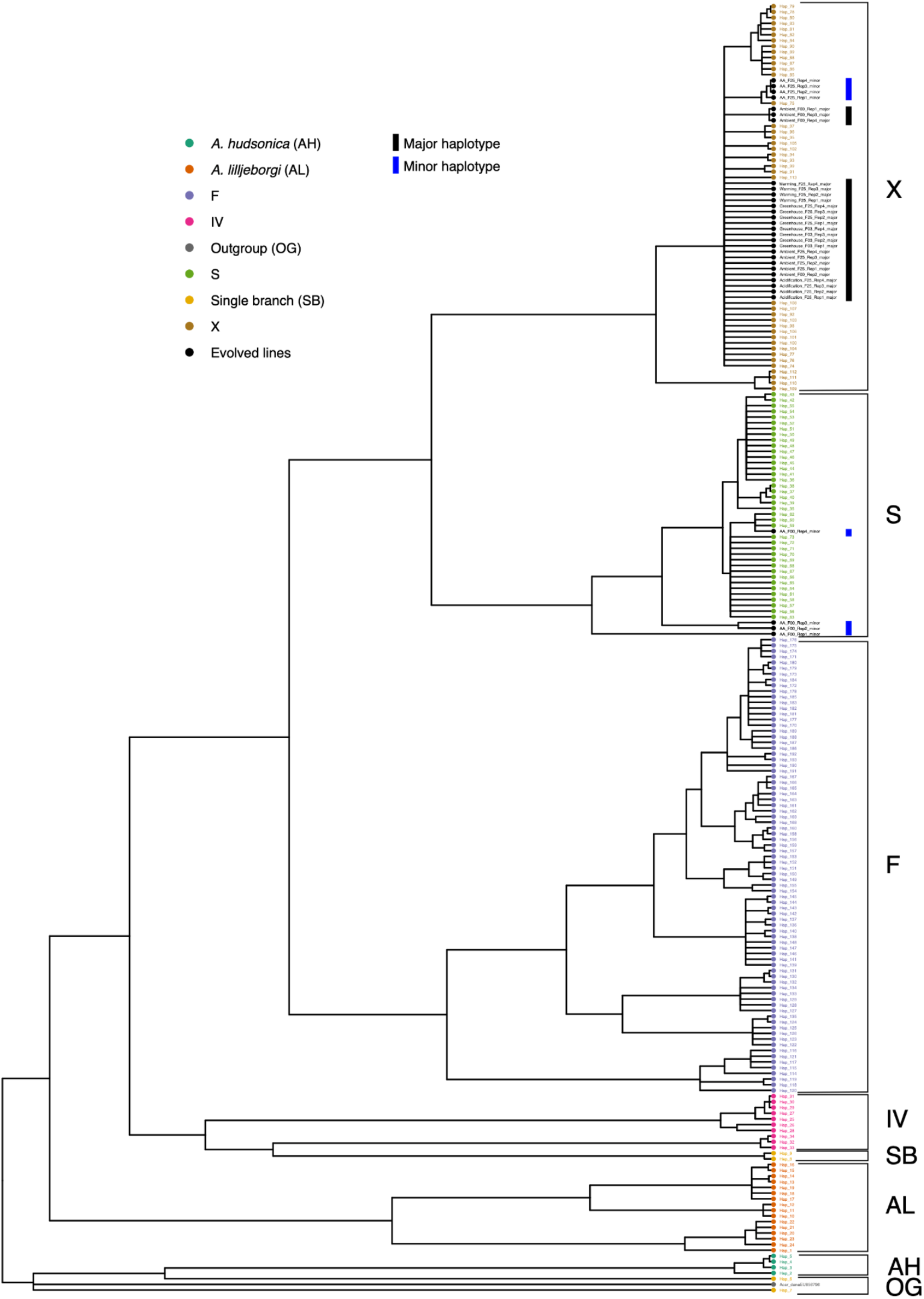
Phylogenetic tree from COI sequence data. We determined the clade of samples from this study using samples from Figueroa et al. (2020). Black and blue bars are the tree tips indicate major and minor haplotypes in the pooled samples, respectively. Samples in our study were predominantly clade X.

**Figure S12:**
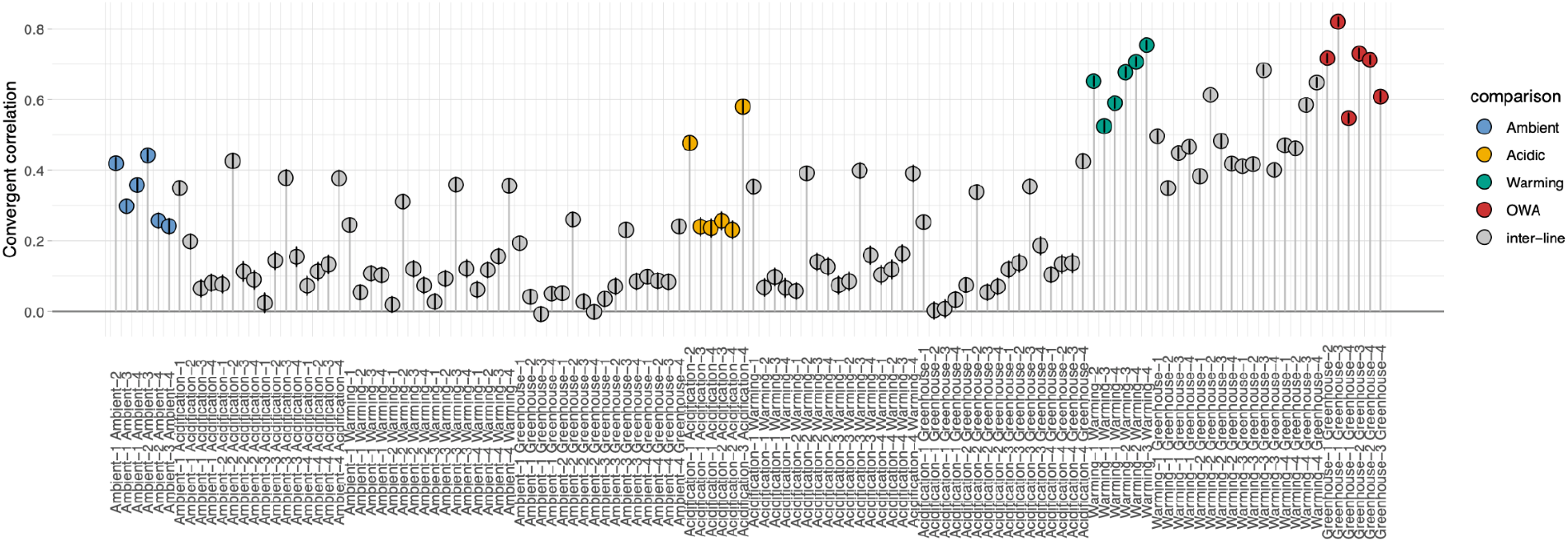
Convergent correlations of allele frequency change from F0 to F25 between samples. Black lines for each point show the 95% bootstrap confidence interval. Note the increase in convergent correlation between samples sharing the same replicate number due to an artefact from calculating allele frequency change from the same F0 sample. Samples with shared F0 samples (i.e., the same replicate number) were dropped from further calculations.

## Notes

### Competing Interest Statement

The authors have declared no competing interest.

## References

Alexa A, Rahnenfuhrer J. 2019. Gene set enrichment analysis with topGO.

Alexa A, Rahnenführer J, Lengauer T. 2006. Improved scoring of functional groups from gene expression data by decorrelating GO graph structure. Bioinformatics 22:1600–1607.

Ali MH, Mungai PT, Schumacker PT. 2006. Stretch-induced phosphorylation of focal adhesion kinase in endothelial cells: role of mitochondrial oxidants. Am J Physiol Lung Cell Mol Physiol 291:L38–45.

Bailey A, De Wit P, Thor P, Browman HI, Bjelland R, Shema S, Fields DM, Runge JA, Thompson C, Hop H. 2017. Regulation of gene expression is associated with tolerance of the Arctic copepod *Calanus glacialis* to CO2-acidified sea water. Ecol Evol 7:7145–7160.

Banse K. 1995. Zooplankton: Pivotal role in the control of ocean production: I. Biomass and production. ICES J Mar Sci 52:265–277.

Barghi N, Hermisson J, Schlötterer C. 2020. Polygenic adaptation: a unifying framework to understand positive selection. Nat Rev Genet 21:769–781.

Barghi N, Tobler R, Nolte V, Jakšić AM, Mallard F, Otte KA, Dolezal M, Taus T, Kofler R, Schlötterer C. 2019. Genetic redundancy fuels polygenic adaptation in *Drosophila*. PLoS Biol 17:e3000128.

Barrett RDH, Schluter D. 2008. Adaptation from standing genetic variation. Trends Ecol Evol 23:38–44.

Bates D, Mächler M, Bolker B, Walker S. 2014. Fitting Linear Mixed-Effects Models using lme4. arXiv [statCO].

Baumann H, Smith EM. 2018. Quantifying metabolically driven pH and oxygen fluctuations in US nearshore habitats at diel to interannual time scales. Estuaries Coasts.

Baumann H, Wallace RB, Tagliaferri T, Gobler CJ. 2015. Large Natural pH, CO2 and O2 Fluctuations in a Temperate Tidal Salt Marsh on Diel, Seasonal, and Interannual Time Scales. Estuaries Coasts 38:220–231.

Belhadj Slimen I, Najar T, Ghram A, Dabbebi H, Ben Mrad M, Abdrabbah M. 2014. Reactive oxygen species, heat stress and oxidative-induced mitochondrial damage. A review. Int J Hyperthermia 30:513–523.

Ben Mahdi MH, Andrieu V, Pasquier C. 2000. Focal adhesion kinase regulation by oxidative stress in different cell types. IUBMB Life 50:291–299.

Bernatchez L. 2016. On the maintenance of genetic variation and adaptation to environmental change: considerations from population genomics in fishes. J Fish Biol 89:2519–2556.

Bindoff NL, Cheung WWL, Kairo JG, Arístegui J, Guinder VA, Hallberg R, Hilmi NJM, Jiao N, Karim MS, Levin L, O’Donoghue S, Purca Cuicapusa SR, Rinkevich B, Suga T, Tagliabue A, Williamson P. 2019. Changing Ocean, Marine Ecosystems, and Dependent Communities In: Pörtner H-O, Roberts DC, Masson-Delmotte V, Zhai P, Tignor M, Poloczanska E, Mintenbeck K, Alegría A, Nicolai M, Okem A, Petzold J, Rama B, Weyer NM, editors. Switzerland: Intergovernmental Panel on Climate Change. pp. 477–587.

Bitter MC, Kapsenberg L, Gattuso J-P, Pfister CA. 2019. Standing genetic variation fuels rapid adaptation to ocean acidification. Nat Commun 10:5821.

Bolger AM, Lohse M, Usadel B. 2014. Trimmomatic: a flexible trimmer for Illumina sequence data. Bioinformatics 30:2114–2120.

Boyd PW, Collins S, Dupont S, Fabricius K, Gattuso J-P, Havenhand J, Hutchins DA, Riebesell U, Rintoul MS, Vichi M, Others. 2018. Experimental strategies to assess the biological ramifications of multiple drivers of global ocean change—a review. Glob Chang Biol 24:2239–2261.

Brennan RS, Garrett AD, Huber KE, Hargarten H, Pespeni MH. 2019. Rare genetic variation and balanced polymorphisms are important for survival in global change conditions. Proceedings of the Royal Society B: Biological Sciences 286:20190943.

Brennan RS, Healy TM, Bryant HJ, Van La M, Schulte PM, Whitehead A. 2018. Integrative Population and Physiological Genomics Reveals Mechanisms of Adaptation in Killifish. Mol Biol Evol 35:2639–2653.

Brennan RS, James AM, Dam HG, Finiguerra M. 2021. Loss and recovery of transcriptional plasticity after long-term adaptation to global change conditions in a marine copepod. bioRxiv.

Buffalo V, Coop G. 2020. Estimating the genome-wide contribution of selection to temporal allele frequency change. Proc Natl Acad Sci U S A 117:20672–20680.

Buffalo V, Coop G. 2019. The Linked Selection Signature of Rapid Adaptation in Temporal Genomic Data. Genetics 213:1007–1045.

Bumpus HC. 1899. The Elimination of the Unfit as Illustrated by the Introduced Sparrow, Passer Domesticus:(a Fourth Contribution to the Study of Variation). Gin.

Caldeira K, Wickett ME. 2003. Oceanography: anthropogenic carbon and ocean pH. Nature 425:365.

Calliari D, Andersen Borg MC, Thor P, Gorokhova E, Tiselius P. 2008. Instantaneous salinity reductions affect the survival and feeding rates of the co-occurring copepods Acartia tonsa Dana and A. clausi Giesbrecht differently. J Exp Mar Bio Ecol 362:18–25.

Camacho C, Coulouris G, Avagyan V, Ma N, Papadopoulos J, Bealer K, Madden TL. 2009. BLAST+: architecture and applications. BMC Bioinformatics 10:421.

Campbell-Staton SC, Cheviron ZA, Rochette N, Catchen J, Losos JB, Edwards SV. 2017. Winter storms drive rapid phenotypic, regulatory, and genomic shifts in the green anole lizard. Science 357:495–498.

Castro JP, Yancoskie MN, Marchini M, Belohlavy S, Hiramatsu L, Kučka M, Beluch WH, Naumann R, Skuplik I, Cobb J, Barton NH, Rolian C, Chan YF. 2019. An integrative genomic analysis of the Longshanks selection experiment for longer limbs in mice. Elife 8. doi:10.7554/eLife.42014

Cervetto G, Gaudy R, Pagano M. 1999. Influence of salinity on the distribution of Acartia tonsa (Copepoda, Calanoida). J Exp Mar Bio Ecol 239:33–45.

Chen G, Hare MP. 2011. Cryptic diversity and comparative phylogeography of the estuarine copepod Acartia tonsa on the US Atlantic coast. Mol Ecol 20:2425–2441.

Chen G, Hare MP. 2008. Cryptic ecological diversification of a planktonic estuarine copepod, Acartia tonsa. Mol Ecol 17:1451–1468.

Chinnery FE, Williams JA. 2004. The influence of temperature and salinity on Acartia (Copepoda: Calanoida) nauplii survival. Mar Biol 145:733–738.

Chung DJ, Schulte PM. 2020. Mitochondria and the thermal limits of ectotherms. J Exp Biol 223. doi:10.1242/jeb.227801

Dam HG, deMayo JA, Park G, Norton L, He X, Finiguerra MB, Baumann H, Brennan RS, Pespeni MH. 2021. Rapid, but limited, zooplankton adaptation to simultaneous warming and acidification. Nat Clim Chang 11:780–786.

Dam HG, Zhang X, Butler M, Roman MR. 1995. Mesozooplankton grazing and metabolism at the equator in the central Pacific: Implications for carbon and nitrogen fluxes. Deep Sea Res Part 2 Top Stud Oceanogr 42:735–756.

de Vries A, Ripley BD. 2016. ggdendro: Create Dendrograms and Tree Diagrams Using’ggplot2’. R package version 0 1-20.

De Wit P, Dupont S, Thor P. 2016. Selection on oxidative phosphorylation and ribosomal structure as a multigenerational response to ocean acidification in the common copepod *Pseudocalanus acuspes*. Evol Appl 9:1112–1123.

De Wit P, Rogers-Bennett L, Kudela RM, Palumbi SR. 2014. Forensic genomics as a novel tool for identifying the causes of mass mortality events. Nat Commun 5:3652.

Doney SC, Fabry VJ, Feely RA, Kleypas JA. 2009. Ocean acidification: the other CO2 problem. Ann Rev Mar Sci 1:169–192.

Donihue CM, Herrel A, Fabre A-C, Kamath A, Geneva AJ, Schoener TW, Kolbe JJ, Losos JB. 2018. Hurricane-induced selection on the morphology of an island lizard. Nature. doi:10.1038/s41586-018-0352-3

Faust O, Abayev-Avraham M, Wentink AS, Maurer M, Nillegoda NB, London N, Bukau B, Rosenzweig R. 2020. HSP40 proteins use class-specific regulation to drive HSP70 functional diversity. Nature 587:489–494.

Feder AF, Petrov DA, Bergland AO. 2012. LDx: estimation of linkage disequilibrium from high-throughput pooled resequencing data. PLoS One 7:e48588.

Feder ME, Hofmann GE. 1999. Heat-shock proteins, molecular chaperones, and the stress response: evolutionary and ecological physiology. Annu Rev Physiol 61:243–282.

Feinberg LR, Dam HG. 1998. Effects of diet on dimensions, density and sinking rates of fecal pellets of the copepod Acartia tonsa. Mar Ecol Prog Ser 175:87–96.

Figueroa NJ, Figueroa DF, Hicks D. 2020. Phylogeography of Acartia tonsa Dana, 1849 (Calanoida: Copepoda) and phylogenetic reconstruction of the genus Acartia Dana, 1846. Mar Biodivers 50:23.

Garrett AD, Brennan RS, Steinhart AL, Pelletier AM, Pespeni MH. 2020. Unique genomic and phenotypic responses to extreme and variable pH conditions in purple urchin larvae. Integr Comp Biol. doi:10.1093/icb/icaa072

Gerber L, Lee CE, Grousset E, Blondeau-Bidet E, Boucheker NB, Lorin-Nebel C, Charmantier-Daures M, Charmantier G. 2016. The legs have it: In situ expression of ion transporters V-type H(+)-ATPase and Na(+)/K(+)-ATPase in the osmoregulatory leg organs of the invading copepod Eurytemora affinis. Physiol Biochem Zool 89:233–250.

Goncalves P, Thompson EL, Raftos DA. 2017. Contrasting impacts of ocean acidification and warming on the molecular responses of CO2-resilient oysters. BMC Genomics 18:431.

Griffiths JS, Kawji Y, Kelly MW. 2021. An Experimental Test of Adaptive Introgression in Locally Adapted Populations of Splash Pool Copepods. Mol Biol Evol 38:1306–1316.

Gunderson AR, Armstrong EJ, Stillman JH. 2016. Multiple Stressors in a Changing World: The Need for an Improved Perspective on Physiological Responses to the Dynamic Marine Environment. Ann Rev Mar Sci 8:357–378.

Haas BJ, Papanicolaou A, Yassour M, Grabherr M, Blood PD, Bowden J, Couger MB, Eccles D, Li B, Lieber M, MacManes MD, Ott M, Orvis J, Pochet N, Strozzi F, Weeks N, Westerman R, William T, Dewey CN, Henschel R, LeDuc RD, Friedman N, Regev A. 2013. De novo transcript sequence reconstruction from RNA-seq using the Trinity platform for reference generation and analysis. Nat Protoc 8:1494–1512.

Harada AE, Healy TM, Burton RS. 2019. Variation in thermal tolerance and its relationship to mitochondrial function across populations of *Tigriopus californicus*. Front Physiol 10:213.

Hermisson J, Pennings PS. 2005. Soft sweeps: molecular population genetics of adaptation from standing genetic variation. Genetics 169:2335–2352.

Höllinger I, Pennings PS, Hermisson J. 2019. Polygenic adaptation: From sweeps to subtle frequency shifts. PLoS Genet 15:e1008035.

Hsu S-K, Belmouaden C, Nolte V, Schlötterer C. 2021. Parallel gene expression evolution in natural and laboratory evolved populations. Mol Ecol 30:884–894.

Jørgensen TS, Petersen B, Petersen HCB, Browne PD, Prost S, Stillman JH, Hansen LH, Hansen BW. 2019. The Genome and mRNA transcriptome of the cosmopolitan calanoid copepod *Acartia tonsa* Dana improve the understanding of copepod genome size evolution. Genome Biol Evol 11:1440–1450.

Kelly JK, Hughes KA. 2019. Pervasive Linked Selection and Intermediate-Frequency Alleles Are Implicated in an Evolve-and-Resequencing Experiment of Drosophila simulans. Genetics 211:943–961.

Kelly MW, Griffiths JS. 2021. Selection Experiments in the Sea: What Can Experimental Evolution Tell Us About How Marine Life Will Respond to Climate Change? Biol Bull 000–000.

Kelly MW, Pankey MS, DeBiasse MB, Plachetzki DC. 2017. Adaptation to heat stress reduces phenotypic and transcriptional plasticity in a marine copepod. Funct Ecol 31:398–406.

Kelly MW, Sanford E, Grosberg RK. 2012. Limited potential for adaptation to climate change in a broadly distributed marine crustacean. Proc Biol Sci 279:349–356.

Kemper KE, Saxton SJ, Bolormaa S, Hayes BJ, Goddard ME. 2014. Selection for complex traits leaves little or no classic signatures of selection. BMC Genomics 15:246.

Koboldt DC, Zhang Q, Larson DE, Shen D, McLellan MD, Lin L, Miller CA, Mardis ER, Ding L, Wilson RK. 2012. VarScan 2: somatic mutation and copy number alteration discovery in cancer by exome sequencing. Genome Res 22:568–576.

Kofler R, Pandey RV, Schlötterer C. 2011. PoPoolation2: identifying differentiation between populations using sequencing of pooled DNA samples (Pool-Seq). Bioinformatics 27:3435–3436.

Kofler R, Schlötterer C. 2014. A guide for the design of evolve and resequencing studies. Mol Biol Evol 31:474–483.

Langer JAF, Meunier CL, Ecker U, Horn HG, Schwenk K, Boersma M. 2019. Acclimation and adaptation of the coastal calanoid copepod *Acartia tonsa* to ocean acidification: a long-term laboratory investigation. Mar Ecol Prog Ser 619:35–51.

Láruson ÁJ, Yeaman S, Lotterhos KE. 2020. The importance of genetic redundancy in evolution. Trends Ecol Evol 35:809–822.

Lee CE, Kiergaard M, Gelembiuk GW, Eads BD, Posavi M. 2011. Pumping ions: rapid parallel evolution of ionic regulation following habitat invasions. Evolution 65:2229–2244.

Lee CE, Remfert JL, Chang Y-M. 2007. Response to selection and evolvability of invasive populations. Genetica 129:179–192.

Lewis CN, Brown KA, Edwards LA, Cooper G, Findlay HS. 2013. Sensitivity to ocean acidification parallels natural pCO2 gradients experienced by Arctic copepods under winter sea ice. Proc Natl Acad Sci U S A 110:E4960–7.

Li H. 2013. Aligning sequence reads, clone sequences and assembly contigs with BWA-MEM. arXiv preprint arXiv:13033997.

Mallard F, Nolte V, Schlötterer C. 2020. The evolution of phenotypic plasticity in response to temperature stress. Genome Biol Evol 12:2429–2440.

Mallard F, Nolte V, Tobler R, Kapun M, Schlötterer C. 2018. A simple genetic basis of adaptation to a novel thermal environment results in complex metabolic rewiring in *Drosophila*. Genome Biol 19:119.

Mayer MP, Bukau B. 2005. Hsp70 chaperones: cellular functions and molecular mechanism. Cell Mol Life Sci 62:670–684.

Melzner F, Gutowska MA, Langenbuch M, Dupont S, Lucassen M, Thorndyke MC, Bleich M, Pörtner H-O. 2009. Physiological basis for high CO2 tolerance in marine ectothermic animals: pre-adaptation through lifestyle and ontogeny? Biogeosci Discuss 6:4693–4738.

Messer PW, Ellner SP, Hairston NG Jr. 2016. Can Population Genetics Adapt to Rapid Evolution? Trends Genet 32:408–418.

Messer PW, Petrov DA. 2013. Population genomics of rapid adaptation by soft selective sweeps. Trends Ecol Evol 28:659–669.

Möllmann C, Müller-Karulis B, Kornilovs G, St John MA. 2008. Effects of climate and overfishing on zooplankton dynamics and ecosystem structure: regime shifts, trophic cascade, and feedback loops in a simple ecosystem. ICES J Mar Sci 65:302–310.

Nixon SW, Granger S, Buckley BA, Lamont M, Rowell B. 2004. A one hundred and seventeen year coastal water temperature record from Woods Hole, Massachusetts. Estuaries 27:397–404.

Pelejero C, Calvo E, Hoegh-Guldberg O. 2010. Paleo-perspectives on ocean acidification. Trends Ecol Evol 25:332–344.

Pennings PS, Hermisson J. 2006. Soft sweeps II--molecular population genetics of adaptation from recurrent mutation or migration. Mol Biol Evol 23:1076–1084.

Pespeni MH, Sanford E, Gaylord B, Hill TM, Hosfelt JD, Jaris HK, LaVigne M, Lenz EA, Russell AD, Young MK, Palumbi SR. 2013. Evolutionary change during experimental ocean acidification. Proc Natl Acad Sci U S A 110:6937–6942.

Plough LV, Fitzgerald C, Plummer A, Pierson JJ. 2018. Reproductive isolation and morphological divergence between cryptic lineages of the copepod Acartia tonsa in Chesapeake Bay. Mar Ecol Prog Ser 597:99–113.

Pritchard JK, Di Rienzo A. 2010. Adaptation - not by sweeps alone. Nat Rev Genet 11:665–667.

Pritchard JK, Pickrell JK, Coop G. 2010. The genetics of human adaptation: hard sweeps, soft sweeps, and polygenic adaptation. Curr Biol 20:R208–15.

Rodríguez-Trelles F, Tarrío R, Santos M. 2013. Genome-wide evolutionary response to a heat wave in Drosophila. Biol Lett 9:20130228.

Ronquist F, Teslenko M, van der Mark P, Ayres DL, Darling A, Höhna S, Larget B, Liu L, Suchard MA, Huelsenbeck JP. 2012. MrBayes 3.2: efficient Bayesian phylogenetic inference and model choice across a large model space. Syst Biol 61:539–542.

Sasaki M, Dam HG. 2021. Global patterns in copepod thermal tolerance. J Plankton Res 43:598–609.

Schlötterer C, Kofler R, Versace E, Tobler R, Franssen SU. 2015. Combining experimental evolution with next-generation sequencing: a powerful tool to study adaptation from standing genetic variation. Heredity 114:431–440.

Schoville SD, Barreto FS, Moy GW, Wolff A, Burton RS. 2012. Investigating the molecular basis of local adaptation to thermal stress: population differences in gene expression across the transcriptome of the copepod Tigriopus californicus. BMC Evol Biol 12:170.

Sørensen JG. 2010. Application of heat shock protein expression for detecting natural adaptation and exposure to stress in natural populations. Curr Zool 56:703–713.

Stern DB, Lee CE. 2020. Evolutionary origins of genomic adaptations in an invasive copepod. Nat Ecol Evol 4:1084–1094.

Storey JD, Base AJ, Dabney A, Robinson D. 2019. qvalue: Q-value estimation for false discovery rate control.

Tangwancharoen S, Moy GW, Burton RS. 2018. Multiple Modes of Adaptation: Regulatory and Structural Evolution in a Small Heat Shock Protein Gene. Mol Biol Evol 35:2110–2119.

Taus T, Futschik A, Schlötterer C. 2017. Quantifying Selection with Pool-Seq Time Series Data. Mol Biol Evol 34:3023–3034.

Therkildsen NO, Wilder AP, Conover DO, Munch SB, Baumann H, Palumbi SR. 2019. Contrasting genomic shifts underlie parallel phenotypic evolution in response to fishing. Science 365:487–490.

Thor P, Dupont S. 2015. Transgenerational effects alleviate severe fecundity loss during ocean acidification in a ubiquitous planktonic copepod. Glob Chang Biol 21:2261–2271.

Tobler R, Franssen SU, Kofler R, Orozco-Terwengel P, Nolte V, Hermisson J, Schlötterer C. 2014. Massive habitat-specific genomic response in *D. melanogaster* populations during experimental evolution in hot and cold environments. Mol Biol Evol 31:364–375.

Turner JT. 1984. The feeding ecology of some zooplankters that are important prey items of larval fish (No. 7). NOAA Technical Report NMFS.

Vargas CA, Lagos NA, Lardies MA, Duarte C, Manríquez PH, Aguilera VM, Broitman B, Widdicombe S, Dupont S. 2017. Species-specific responses to ocean acidification should account for local adaptation and adaptive plasticity. Nat Ecol Evol 1:84.

Vlachos C, Burny C, Pelizzola M, Borges R, Futschik A, Kofler R, Schlötterer C. 2019. Benchmarking software tools for detecting and quantifying selection in evolve and resequencing studies. Genome Biol 20:169.

Waldvogel A-M, Feldmeyer B, Rolshausen G, Exposito-Alonso M, Rellstab C, Kofler R, Mock T, Schmid K, Schmitt I, Bataillon T, Savolainen O, Bergland A, Flatt T, Guillaume F, Pfenninger M. 2020. Evolutionary genomics can improve prediction of species’ responses to climate change. Evol Lett 4:4–18.

Wang M, Zhao Y, Zhang B. 2015. Efficient Test and Visualization of Multi-Set Intersections. Sci Rep 5:16923.

Whiteley NM. 2011. Physiological and ecological responses of crustaceans to ocean acidification. Mar Ecol Prog Ser 430:257–271.

Wu TD, Watanabe CK. 2005. GMAP: a genomic mapping and alignment program for mRNA and EST sequences. Bioinformatics 21:1859–1875.

Yu G, Li F, Qin Y, Bo X, Wu Y, Wang S. 2010. GOSemSim: an R package for measuring semantic similarity among GO terms and gene products. Bioinformatics 26:976–978.

## Supplemental References

Edgar RC. 2004. MUSCLE: a multiple sequence alignment method with reduced time and space complexity. BMC Bioinformatics 5:113.

Paradis E, Claude J, Strimmer K. 2004. APE: Analyses of Phylogenetics and Evolution in R language. Bioinformatics 20:289–290.

Yu G, Smith DK, Zhu H, Guan Y, Lam TT. 2017. Ggtree: An r package for visualization and annotation of phylogenetic trees with their covariates and other associated data. Methods Ecol Evol 8:28–36.

